# Reactivation of the progenitor gene Trim71 enhances the mitotic and hair cell-forming potential of cochlear supporting cells

**DOI:** 10.1101/2023.01.12.523802

**Authors:** Xiao-Jun Li, Charles Morgan, Prathamesh T Nadar-Ponniah, Waldemar Kolanus, Angelika Doetzlhofer

**Author notes:** **Correspondence addressed to:** Angelika Doetzlhofer, Ph.D., The Solomon H. Snyder Department of Neuroscience, Johns Hopkins School of Medicine, 855 N. Wolfe Street, Rangos Building, Room 433, Baltimore, MD 21205.

## Abstract

Cochlear hair cell loss is a leading cause of deafness in humans. Neighboring supporting cells have some capacity to regenerate hair cells. However, their regenerative potential sharply declines as supporting cells undergo maturation (postnatal day 5 in mice). We recently reported that reactivation of the RNA-binding protein LIN28B restores the hair cell-regenerative potential of P5 cochlear supporting cells. Here, we identify the LIN28B target *Trim71* as a novel and equally potent enhancer of supporting cell plasticity. TRIM71 is a critical regulator of stem cell behavior and cell reprogramming, however, its role in cell regeneration is poorly understood. Employing an organoid-based assay, we show that TRIM71 reactivation increases the mitotic and hair cell-forming potential of P5 cochlear supporting cells by facilitating their de-differentiation into progenitor-like cells. Our mechanistic work indicates that TRIM71’s RNA-binding activity is essential for such ability, and our transcriptomic analysis identifies gene modules that are linked to TRIM71 and LIN28B-mediated supporting cell reprogramming. Furthermore, our study uncovers that the TRIM71-LIN28B target *Hmga2* is essential for supporting cell self-renewal and hair cell formation.

## Introduction

Mechano-sensory hair cells within the auditory sensory organ (termed cochlea) are essential for our perception of sound. In mammals, auditory hair cells do not regenerate, and their loss or dysfunction is a leading cause for deafness in humans. In non-mammalian vertebrates, such as birds and fish, damaged or lost hair cells are replenished by their surrounding supporting cells through mitotic and non-mitotic mechanisms (reviewed in (Brignull et al., 2009)). A recent analysis of hair cell regeneration in the zebrafish lateral line revealed that once initiated, hair cell regeneration recapitulates hair cell development (Baek et al., 2022). Hair cells and supporting cells in the vertebrate inner ear originate from SOX2 and Jagged1 (JAG1) expressing embryonic progenitor cells termed pro-sensory cells (Kiernan et al., 2005, Gu et al., 2016). Hair cell fate is determined by the transcription factor ATOH1, which is both necessary and sufficient for hair cell formation (Bermingham et al., 1999, Zheng and Gao, 2000). *Atoh1* induction in pro-sensory cells is controlled, amongst others, by Notch-mediated lateral inhibition and Wnt/-β-catenin signaling. Activation of canonical Wnt signaling promotes cell mitosis and activates *Atoh1* expression (Jacques et al., 2012, Shi et al., 2014), while high levels of Notch signaling inhibits the induction of *Atoh1* expression, and forces pro-sensory cells to adopt a supporting cell fate (Takebayashi et al., 2007). In mice, cochlear hair cells form between embryonic days 14.5 (E14.5) and E18.5. However, many developmental gene programs engaged during hair cell formation are still active after birth. For instance, in the presence of mitogens and cultured on a mesenchymal feeder layer, cochlear supporting cells, isolated from early postnatal mice [postnatal day 0 (P0)-P3] readily re-enter the cell cycle, proliferate and form hair cells (White et al., 2006, Sinkkonen et al., 2011). Hair cell formation by cochlear supporting cells is also induced by inhibition of Notch signaling (Mizutari et al., 2013, Korrapati et al., 2013) or over-activation of Wnt/-β-catenin signaling, which at early postnatal stages also promotes cell cycle re-entry of cochlear supporting cells (Chai et al., 2012, Shi et al., 2012). An even higher potential for mitotic hair cell formation has been demonstrated for Kölliker’s cells, which are a transient population of supporting cell-like epithelial cells that reside at the medial border of the sensory epithelium (Sinkkonen et al., 2011, Kubota et al., 2021). However, the regenerative potential of cochlear supporting cells and surrounding non-sensory epithelial cells wanes during the first postnatal week and once mice are hearing, around postnatal day 12 (P12), ectopic activation of *Atoh1* expression or activation of hair cell-fate inducing signals have thus far failed to generate/regenerate functional hair cells (reviewed in (Atkinson et al., 2015)). We recently uncovered a central role for *let-7* miRNAs and their mutual antagonist LIN28B in the developmental decline in supporting cells plasticity. LIN28B functions as a RNA-binding protein and in the developing cochlea is highly expressed in pro-sensory cells, where it promotes a proliferative, undifferentiated state, while differentiating hair cells and supporting cells express members of the *let-7* family of miRNAs, which act to reinforce a post-mitotic terminal differentiated state (Golden et al., 2015). We found that expression of human *LIN28B* in early postnatal cochlear tissue/cells extends the window of hair cell-regenerative capacity, while overexpression of *let-7g* or loss of *Lin28b* (and *Lin28a*) accelerates the developmental decline of regenerative potential (Li and Doetzlhofer, 2020). Our most recent findings indicate that LIN28B enhances the regenerative capacity of cochlear supporting cells and Kölliker’s cells by reprogramming them into progenitor-like cells. Among the genes that were induced by LIN28B overexpression was *Trim71* (Li et al., 2022). In the developing cochlear epithelial duct *Trim71* expression is limited to undifferentiated progenitor cells (Golden et al., 2015). In general, *Trim71* gene is highly expressed during early embryonic development and functions to promote cell reprogramming and stemness through its dual role as an ubiquitin ligase and RNA binding protein (Worringer et al., 2014, Torres-Fernandez et al., 2021, Duy et al., 2022). *Lin28b* and *Trim71* are evolutionary conserved targets of *let-7* mediated gene silencing (Reinhart et al., 2000, Lin et al., 2007). Recent studies suggest that like LIN28B, TRIM71 interferes with the processing and activity of *let-7* miRNAs (Liu et al., 2021, Torres Fernandez et al., 2021, Rybak et al., 2009). Whether reactivation of *Trim71* enhances the regenerative capacity of cochlear supporting cells has yet to be determined. In this study, we used a cochlear organoid platform as a model to investigate the role of *Trim71* in cochlear supporting cell plasticity. We show that the functionally-related proteins TRIM71 and LIN28B are equally potent enhancers of cochlear supporting cell plasticity. Our mechanistic studies indicate that similar to LIN28B, TRIM71 increase the mitotic and regenerative potential of cochlear supporting cells (and Kölliker’s cells) by promoting their de-differentiation into progenitor-like cells and that such activity is independent of TRIM71’s function as an ubiquitin ligase. Furthermore, our transcriptomic data analysis identifies shared and unique target genes among TRIM71 and LIN28B and our functional analysis of LIN28B and TRIM71 target gene *Hmga2* reveals that *Hmga2* is essential for the ability of cochlear supporting cells/Kölliker’s cells to re-enter the cell cycle and form hair cells.

## Results

### TRIM71 increases the mitotic and hair cell-forming potential of cochlear supporting cells

Cochlear supporting cells are post-mitotic cells, and their re-entry into the cell cycle is an important aspect of the hair cell regenerative process. To determine whether TRIM71 promotes cell cycle reentry of cochlear supporting cells (and Kölliker’s cells), we expressed human *TRIM71* in cochlear organoid cultures that have been established with cochlear epithelial cells obtained from 5-day old mice (postnatal day 5 (P5)). The stage was chosen as at stage P5 cochlear epithelial cells/supporting cells do still re-enter the cell cycle and proliferate but do so at a significant lower frequency than observed at earlier stages (e.g. P2) when placed in organoid culture (Li and Doetzlhofer, 2020, Li et al., 2022). Briefly, we infected cochlear epithelial cells with lentiviral particles that expressed full-length TRIM71 (TRIM71) or mutant forms of TRIM71 protein that lacked the RING (ΔRING), Coiled-Coil (ΔCoiled-Coil) or NHL domain (ΔNHL) (Figure S1A). The RING domain located within the N-terminal tripartite motif is critical for TRIM71’s ubiquitin ligase activity (Rybak et al., 2009), while the coiled-coil domain and C-terminal NHL domain are important hubs for protein interactions and are essential for TRIM71’s RNA binding activity (Loedige et al., 2013). To be able to track infected cells, mCherry was co-expressed, and to be able to confirm TRIM71 expression the full-length and mutant TRIM71 proteins contained an N-terminal HA-tag (Figure S1B-D). As control, cochlear epithelial cells were infected with lentiviral particles that only expressed HA and mCherry. The infected cells were then placed at high density into a Matrigel matrix and cultured in the presence of the growth factors EGF and FGF2, GSK3-β inhibitor CHIR99021, TGFBRI inhibitor 616452 and HDAC inhibitor valproic acid (VPA) for 10 days (Figure 1A). We found that TRIM71 expression increased organoid formation efficiency by 1.5-fold compared to control (Figure 1B), and increased the average organoid diameter by 2-fold compared to control (Figure 1C). Expression of TRIM71 protein that lacked the RING domain (ΔRING), had a similar positive effect as expression of full length TRIM71 protein, indicating that TRIM71 does not require its E3 ubiquitin ligase activity to promote organoid formation and growth (Figures 1B,C). By contrast, expression of TRIM71 protein that lacked the NHL domain (ΔNHL) failed to increase organoid formation or organoid size compared to control (Figures 1B,C), indicating that the RNA binding domain is required for TRIM71’s positive effect on organoid formation and growth. An intermediate phenotype was observed in cultures that expressed Coiled-Coil domain deficient TRIM71 protein (ΔCoiled-Coil), with no defects in organoid formation but a mild reduction in organoid growth (Figures 1B, C). To determine whether *TRIM71* expression enhanced supporting cell/Kölliker’s cell proliferation, we pulsed control and *TRIM71*-expressing cultures with EdU and analyzed the percentage of infected cells per organoid that incorporated EdU. To verify supporting cells/Kölliker’s cell identity, cultures were also stained for Notch ligand Jagged1 (JAG1), which is highly expressed in supporting cells/Kölliker’s cells and their progenitors. We found that more than 95% of cells in control and *TRIM71-*expressing cultures expressed JAG1 (Figure 1D), confirming that the vast majority of organoids are composed of supporting cells/Kölliker’s cells or their precursors. As anticipated, we found that the percentage of EdU^+^ mCherry^+^ cells in *TRIM71-*expressing organoids was 2-fold increased (Figures 1D, E), indicating TRIM71 as a positive regulator of supporting cell/Kölliker’s cell proliferation.

**Figure 1:**
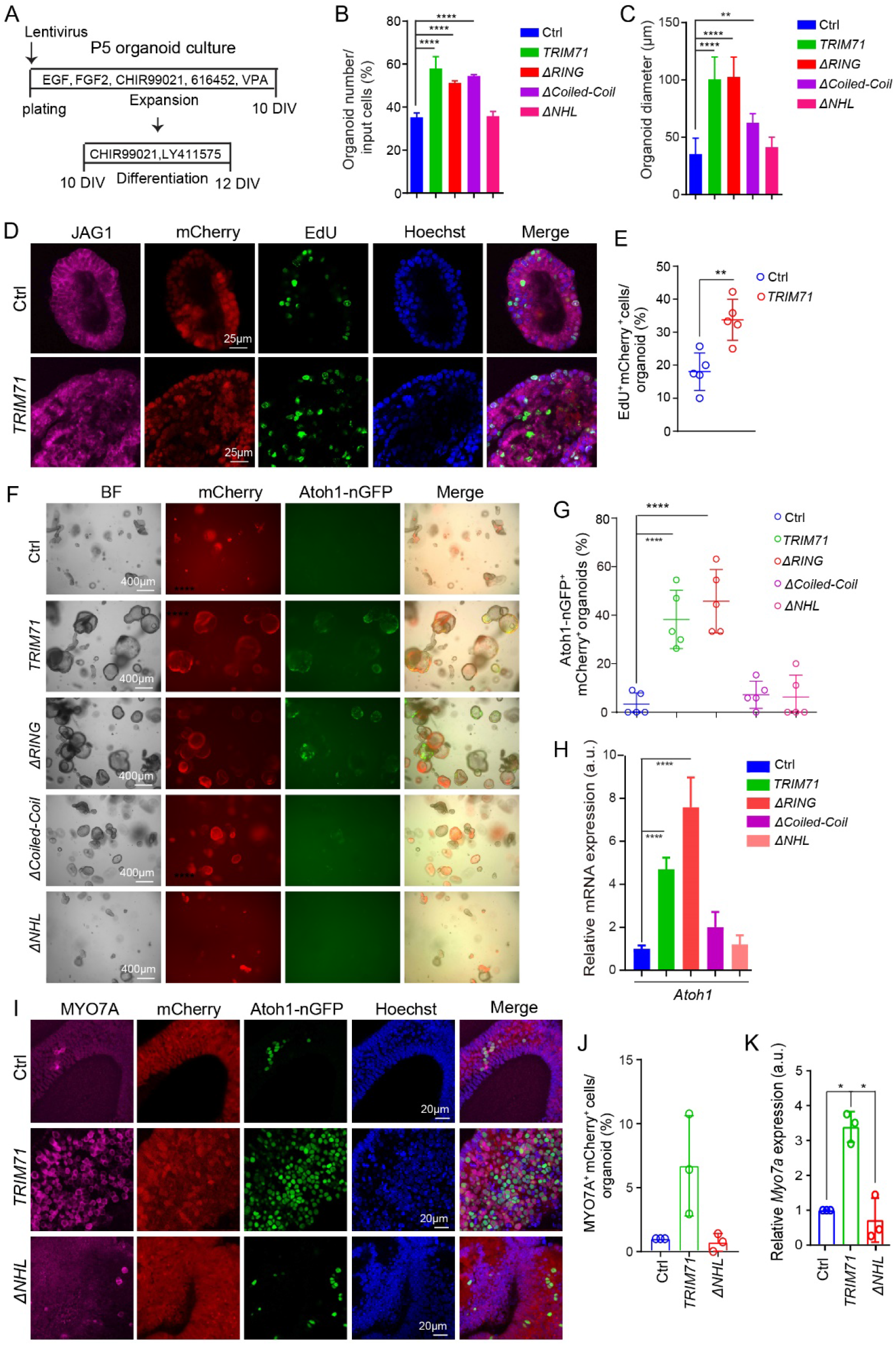
*TRIM71* expression increases the mitotic and hair cell-forming potential of cochlear supporting cells/Kölliker’s cells. **(A)** Experimental scheme. Cochlear epithelial cells from P5 Atoh1-BFP transgenic mice were infected with mCherry expressing control (Ctrl) virus, or virus that co-expressed mCherry and full-length (TRIM71) or mutant TRIM71 protein (Δ RING, Δ Coiled Coil, Δ NHL) **(B-C)** Colony forming efficiency (B) (n=4, three independent experiments) and organoid diameter in (C) (n=4, three independent experiments) after 10 days of expansion. (**D**) Cell proliferation in control and *TRIM71*-expressing organoids. An EdU pulse was given at day 8 and EdU incorporation (red) was analyzed 1.5 hours later. JAG1 (green) marks supporting cells/prosensory cells, mCherry (red) marks infected cells, Hoechst labels cell nuclei (blue). (**E**) Percentage of EdU^+^ mCherry^+^ cells per organoids in (D) (n = 5, two independent experiments). **(F)** Bright field (BF), red (mCherry) and green (Atoh1-nGFP) fluorescent images of organoid cultures. **(C)** Percentage of Atoh1-nGFP^+^mCherry^+^ organoids in (B) (n=5, three independent experiments). **(D)** RT-qPCR of *Atoh1* mRNA in organoids (n=3, two independent experiments). **(E-G)** Confocal images of MYO7A immuno-stained (magenta) organoids Atoh1-GFP (green) and MYO7A (magenta) marks nascent hair cells. MCherry (red) marks infected cells. **(F)** Quantification of MYO7A^+^ mCherry^+^ cells per organoid in (E). **(G)** RT-PCR of *Myo7a* mRNA expression in organoids. Individual data points represent the average value per animal. P-values were calculated using one-way ANOVA with Tukey’s correction with the exception of (E) were two-tailed, unpaired *t* test was used. \**P* ≤ 0.05, \*\**P* < 0.01, \*\*\**P* < 0.001 and \*\*\*\**P* < 0.0001.

We next examined whether expression of *TRIM71* enhances the hair cell-forming capacity of stage P5 cochlear supporting cells/Kölliker’s cells. At stage P5, cochlear epithelial cells (supporting cells and Kölliker’s cells) still produce some hair cells in organoid culture but the rate of hair cell production is greatly reduced compared to earlier stages (e.g. P2) (Li and Doetzlhofer, 2020, Li et al., 2022). To be able to monitor the dynamics of hair cell formation in our cultures, we used cochlear epithelia cells from *Atoh1-nGFP* transgenic mice, which express nuclear GFP in nascent hair cells. To stimulate hair cell formation, organoids were cultured in a ‘differentiation media’ that contained GSK-3β inhibitor CHIR99021 (activates Wnt signaling) and γ-secretase inhibitor LY411575 (inhibits Notch signaling) (Figure 1A). We found that after 2 days of differentiation about 40% of *TRIM71-expressing* organoids contained clusters of nascent hair cells (Atoh1-GFP^+^ cells), while only about 5% of control organoids contained nascent hair cells (Figures 1F, G). Expression of TRIM71 protein that lacked the RING domain (ΔRING) was equally effective in promoting hair cell formation as full-length TRIM71 protein. By contrast TRIM71 protein that lacked either the Coiled-Coil domain (ΔCoiled-Coil) or the NHL (ΔNHL) domain was ineffective in inducing hair cell formation (Figures 1F, G). The positive effect of TRIM71 on hair cell formation was confirmed by analyzing *Atoh1* mRNA expression using RT-qPCR. We found that expression of full-length or RING domain deficient TRIM71 protein resulted in a 4 to 8-fold increase in *Atoh1* mRNA abundance compared to control, while *Atoh1* mRNA levels remained unchanged by the expression of NHL or Coiled-Coil domain deficient TRIM71 protein compared to control (Figure 1H). The hair cell identity of Atoh1-nGFP^+^ cells in *TRIM71-* expressing organoids was confirmed using immunostaining against the hair cell-specific protein myosinVIIa (MYO7A) (Figures 1I, J). Furthermore, we found that TRIM71 expression increased *Myo7a* mRNA expression by more than 3-fold compared to control, while expression of TRIM71 protein that lacked the NHL domain failed to increase *Myo7a* expression (Figure 1K). In summary, our results indicate that expression of TRIM71 protein increases the regenerative capacity of cochlear supporting cells and Kölliker’s cells, and that such activity requires the presence of TRIM71’s NHL or coiled coil domain, whereas TRIM71’s RING-domain is dispensable.

*Trim71* is abundantly expressed in undifferentiated cochlear epithelial cells but its expression rapidly declines at the onset of differentiation and is near undetectable after birth (Golden et al., 2015, Evsen et al., 2020, Kolla et al., 2020). To determine whether *Trim71* is essential for cochlear supporting cells to re-enter the cell cycle and form new hair cells, we deleted *Trim71* in P2 cochlear organoid cultures. We chose stage P2 as at this stage wild-type cochlear epithelial cells, as well as FACS-purified cochlear supporting cells/Kölliker’s cells, readily form large hair cell-containing organoids (Kubota et al., 2021, McLean et al., 2017, Li et al., 2022, Li and Doetzlhofer, 2020). To knockout (KO) *Trim71*, we used *Trim71* floxed (*Trim71^f/f^* mice (Mitschka et al., 2015) that carried *R2^rtTA*M2^* and *TetO-Cre* transgenes for doxycycline-mediated Cre induction (Figure 2A). As control, we used littermates that lacked the *TetO-Cre* transgene. To induce *Trim71* deletion we expanded *TetO-Cre; R26^rtTA*M2^; Trim71^f/f^* (subsequently referred to as *Trim71* KO) and *Trim71^f/f^; R26^rtTA*M2^*(control, wild type) cochlear epithelial cells in the presence of doxycycline (dox) (Figure 2B). After 6 days of expansion, we analyzed organoid formation efficiency and organoid size in control and *Trim71* KO cultures. Our analysis revealed no defects in organoid formation or growth in the absence of *Trim71* (Figures 2C-E). Furthermore, 1.5-hour EdU pulse at 7 days of expansion revealed no significant changes in the rate of cell proliferation in *Trim71* KO organoids compared to control (Figures 2F, G). By contrast, organoid cultures established with cochlear epithelial cells isolated from E13.5 *Trim71* KO embryos formed significantly fewer and smaller organoids (Figures EV2B-D) and incorporated EdU at a significantly lower rate than control cultures established from wild type littermates (Figures EV2E,F). These results indicate that *Trim71* function is essential for cell cycle re-entry and proliferation of undifferentiated cochlear epithelial cells (pro-sensory cells) but its function is not required for the cell cycle reentry and proliferation of cochlear supporting cells (and Kölliker’s cells).

**Figure 2.**
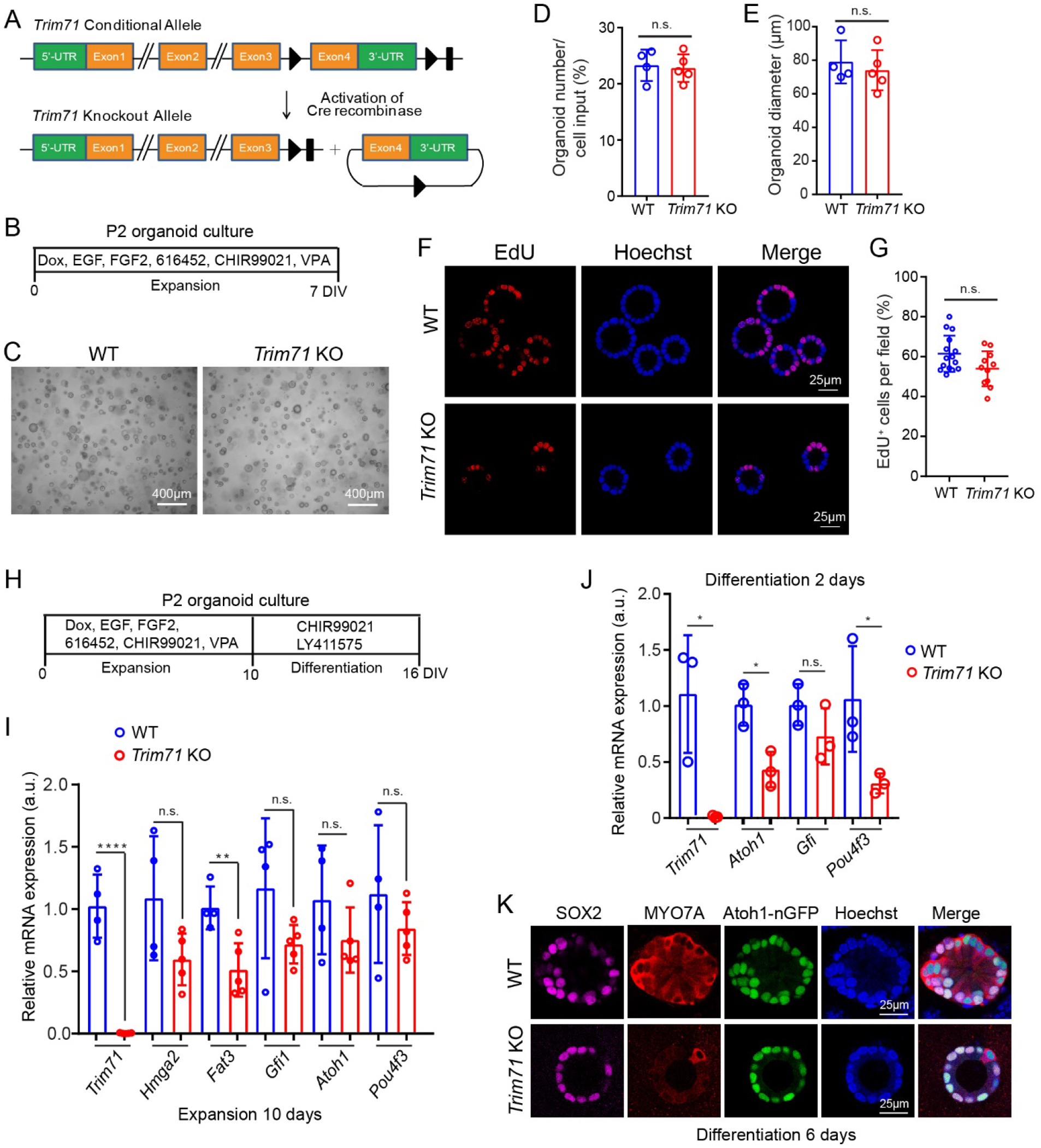
Loss of *Trim71* attenuates the hair cell-forming capacity of cochlear supporting cells/Kölliker’s cells. **(A)** Schematic of *Trim71* conditional and knockout allele. **(B)** Experimental scheme. Cochlear epithelial cells from stage P2 *TetO-Cre;R26 ^rtTA*M2^;Trim71^f/f^* mice (*Trim71* KO) and littermates that lacked *TetO-Cre* transgene (WT) were cultured in the presence of doxycycline (dox). **(C)** Bright field (BF) images of organoid culture after 6 days of expansion. **(D)** Colony forming efficiency in (C). **(E)** Organoid diameter in (C) (n=4 in WT, n=5 in *Trim71* KO, two independent experiments). **(F)** Confocal images of EdU (red) and Hoechst (blue) labeled organoids. A single EdU pulse was given at day 7 and EdU incorporation was analyzed 1.5 hours later. **(G)** Percentage of EdU^+^ cells in (F) (n=15 in WT, n=11 in *Trim71* KO, three independent experiments). **(H)** Experimental scheme. **(I)** RT-qPCR of *Trim71* and progenitor (*Hmag2 and Fat3*) and hair cell-specific (*Atoh1, Gfi1* and *Pou4f3*) mRNAs in control and *Trim71* KO organoids at day 10 (n=4 in WT, n=5 in *Trim71* KO, two independent experiments). **(J)** RT-qPCR of *Trim71* and hair cell-specific (*Atoh1, Gfi1* and *Pou4f3*) mRNAs in WT and *Trim71* KO organoids after 2 days of differentiation (n=3, two independent experiments). **(K)** Confocal images of MYO7A and SOX2 immuno-stained organoids after 6 days of differentiation. Nascent hair cells co-express Atoh1-nGFP (green), MYO7A (red) and SOX2 (magenta). Two-tailed, unpaired *t* test was used to calculate *P* values. \**P* ≤ 0.05, \*\**P* < 0.01 and \*\*\*\**P* < 0.0001.

### Loss of *Trim71* attenuates the hair cell-forming capacity of cochlear supporting cells/Kölliker’s cells

We next analyzed whether loss of *Trim71* impacts the hair cell-forming capacity of cochlear progenitor cells and supporting cells/Kölliker’s cells. Our analysis revealed that while organoid cultures established with E13.5 wild type cochlear epithelial cells readily produced Atoh1-nGFP^+^ hair cells, organoid cultures established with E13.5 *Trim71* deficient cochlear epithelial cells failed to produce Atoh1-nGFP^+^ hair cells (Figures EV2G, H). Much milder defects in hair cell formation were observed in *Trim71* KO organoid cultures established with P2 cochlear epithelial cells. We found that after 2 days of differentiation, P2 *Trim71* KO organoids expressed hair cell-fate inducing genes (*Atoh1, Pou4f3*) on average at a 2-fold lower level than control organoids (Figure 3J). After 6 days of differentiation the majority of Atoh1-nGFP^+^ cells in control organoids co-expressed hair cell-specific protein MYO7A. By contrast, Atoh1-nGFP^+^ cells in *Trim71* KO organoids largely lacked MYO7A expression (Figure 2K). We observed qualitative similar defects in hair cell formation when we used Cre and mCherry expressing lentiviral particles to delete *Trim71* in stage P2 cochlear organoid culture (Figure S2A, B). RT-qPCR revealed that hair cell-specific genes (*Myo7a, Pou4f3*) were expressed at a significantly lower level in *Trim71* KO organoids than control organoids (Figure S2D). To determine whether the defects in hair cell formation were due to defects in supporting cell reprogramming or due to defects in hair cell-fate induction we analyzed the expression of progenitor, supporting cell and hair cell-specific genes in P2 control and *Trim71* KO cochlear organoids using RT-qPCR. Our analysis revealed that *Trim71* KO organoids expressed progenitor-specific genes (*Hmga2, Fat3, Ccnd2, Fst*) at a lower level and supporting cell-specific genes (*Ano1, Cybrd1, S100a1*) at a higher level than control organoids (Figure 2I) (Figure S2C), while the initial induction of hair cell-fate inducing genes (*Atoh1, Pou4f3, Gfi1*) was unaffected (Figure 2I). In sum, our results indicate that prior differentiation *Trim71* is essential for cochlear pro-sensory cells to self-renew and form hair cells and at later stages *Trim71* is dispensable for self-renewal but plays a positive role in the reprogramming of supporting cells into progenitor-like cells and the subsequent formation of hair cells.

**Figure 3.**
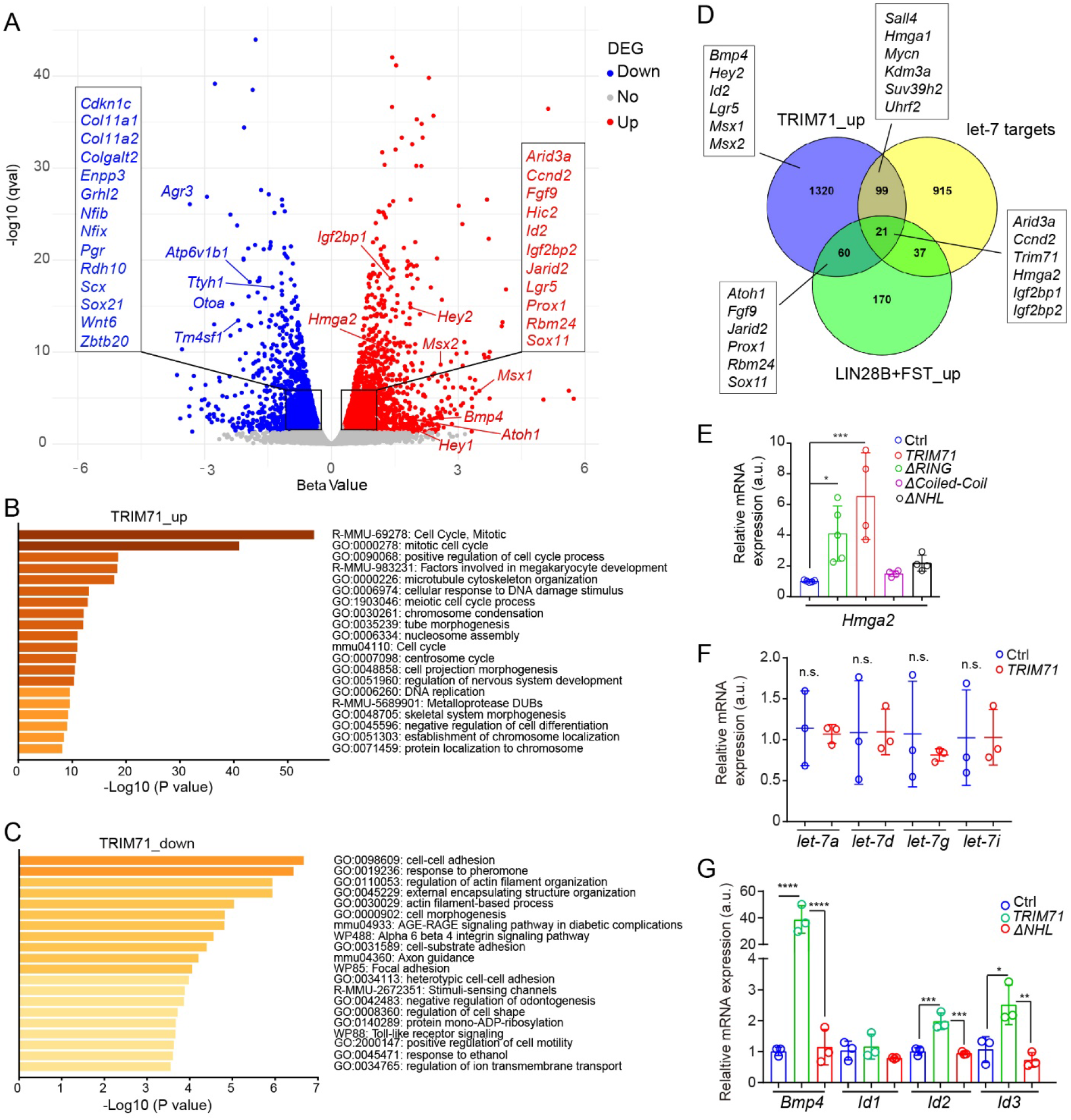
*TRIM71* activates genes linked to self-renewal and de-differentiation in cochlear supporting cells/Kölliker’s’ cells. **(A-C)** Bulk RNA sequencing was used to analyze gene expression in P5 control and *TRIM71*-expressing cochlear organoids at 10 days of expansion. (**A**) Volcano plot of RNA-seq data. Plotted is beta-value (x-axis) versus −log10 q-value (y-axis). Transcripts that are significantly upregulated in response to *TRIM71* expression are marked in red circles, and transcripts that are significantly downregulated are marked in blue circles. (**B**) Biological processes and pathways associated with TRIM71-upregulated genes ranked by adjusted p-value (q-value). (**C**) Biological processes and pathways associated with TRIM71-downregulated genes ranked by adjusted p-value (q-value). (**D**) Venn diagram showing intersection between TRIM71-upregulated genes, predicted *let-7* target genes and LIN28B+FST-upregulated genes. **(E)** RT-qPCR of *Hmga2* mRNA expression in P5 control (Ctrl) cochlear organoids and cochlear organoids that expressed full length (*TRIM71*), RING deficient (Δ*RING*), Coiled-Coil deficient (*ΔCoiled-Coil*) or NHL deficie n t (*ΔNHL*) TRIM71 protein at 10 days of expansion (n=3, two independent experiments). **(F)** TaqMan assay of mature *let-7a-5p, let-7d-5p, let-7g-5p, let-7i-5p* transcripts in P5 control (Ctrl) and *TRIM71* expressing cochlear organoids (n=3, two independent experiments). **(G**) RT-qPCR-based analysis of *Bmp4, Id1, Id2, Id3* expression (n=3, two independent experiments) in control, TRIM71 or TRIM71 ΔNHL expressing cochlear organoids after 10 days of expansion (n=3, two independent experiments). Individual data points represent the average value per animal. One-way ANOVA with Tukey’s correction was used to calculate *P* values. \**P* ≤ 0.05, \*\**P* < 0.01, \*\*\**P* < 0.001 and \*\*\*\**P* < 0.0001.

### TRIM71 increases the expression of genes linked to self-renewal and de-differentiation in cochlear supporting cells/Kölliker’s cells

To uncover the underling mechanism through which TRIM71 enhances the mitotic and hair cell regenerative capacity of supporting cells/Kölliker’s cells, we analyzed the transcriptome of control and T*RIM71*-expressing stage P5 cochlear organoids after 10 days of expansion using RNA-sequencing (RNA-seq). To quantify transcript abundance, we pseudo-align reads to the reference mouse transcriptome (Ensembl Mus musculus v96), using kallisto (v0.46.1) (Bray et al., 2016). We used the companion tool sleuth to determine differentially expressed genes (DEGs) comparing control to *TRIM71*-expressing condition (Pimentel et al., 2017). Our analysis identified 2883 DEGs (q-value< 0.01) (table S1), with 1500 upregulated and 1383 down-regulated genes in response to *TRIM71* expression. Among the TRIM71-upregulated genes were genes that function in pro-sensory cell specification and maintenance (*Hey1, Hey2, Bmp4, Id2, Sox11, Fgf9*) as well as genes that are induced during hair cell formation (*Atoh1, Prox1, Rbm24*) (Figure 3A, red dots). By contrast, the list of TRIM71-downregulated genes included genes critical for cell adhesion (*Chd1, Col11a1, Col11a2*) and glial cell differentiation (*Zbtb20, Nfib, Nfix, Sox21*), including *Nfix*, which transcript has been recently shown to be directly bound by TRIM71 (Foster et al., 2020) (Figure 3A, blue dots). To identify biological processes and pathways that may be altered by *TRIM71* expression, we performed a gene ontology enrichment analysis using Metascape, a web-based portal (Zhou et al., 2019). Consistent with TRIM71’s role as a stemness and pluripotency factor, we found that the list of TRIM71-upregulated genes was significantly enriched for genes that promote cell cycle reentry and mitosis, regulate the response to DNA damage and inhibit cell differentiation (Figure 3B) (table S2), while the list of TRIM71-downregulated genes was significantly enriched for genes that function in cell adhesion, external structure organization as well as actin filament-based processes and organization (Figure 3C)(table S2).

### TRIM71 regulates gene expression in a *let-7* independent manner

LIN28B and TRIM71 proteins are known to enhance each other’s expression and to share many of the same downstream targets (Robinton et al., 2019, Foster et al., 2020). To identify common targets of TRIM71 and LIN28B in supporting cells/Kölliker’s cell, we compared the RNA-seq data with our recently published RNA-seq data that profiled the effects of LIN28B on supporting cell/Kölliker’s cell gene expression in P5 cochlear organoid culture in the presence of follistatin (FST), which we used in lieu of a TGFBR inhibitor (Li et al., 2022). The data comparison revealed that close to one third of LIN28B+FST-upregulated or downregulated genes were in similar fashion up or downregulated by *TRIM71* expression (table S3). The shared list of upregulated genes included well-known *let-7* targets such as *Hmga2, Arid3a, Igf2bp1, Ccnd2* and *Trim71* itself (table S3) (Figure 3D), suggesting that in cochlear organoid culture TRIM71 may inhibit the expression or activity of mature *let-7* miRNAs or both. Recent studies found that TRIM71 represses mature *let-7* miRNA expression by binding LIN28A or LIN28B proteins and enhancing their inhibition of *let-7* biogenesis (Torres Fernandez et al., 2021). Our pull-down experiments using HEK293T cells revealed that NHL or Coiled-Coil deficient TRIM71 proteins failed to bind LIN28B (Figure S3), which correlated with the inability of NHL or Coiled-Coil deficient TRIM71 to upregulate the expression of the *let-7* target *Hmga2* in cochlear organoids (Figure 3E). However, TaqMan-based analysis of mature *let-7a, let-7d, let-7g, let-7i* miRNAs expression in control and TRIM71-expressing organoids revealed no significant changes (Figure 3F). By contrast mature *let-7a, let-7d, let-7g* and *let-7i* miRNAs were near absent in organoids in which we overexpressed human *LIN28B* using a doxycycline-inducible transgenic mouse model (Figure EV3A). TRIM71 destabilizes and inactivates *let-7* miRNAs by inhibiting the expression of Argonaute 2 (AGO2) (Rybak et al., 2009) (Liu et al., 2021). AGO2 is a critical component of the RNA-Induced Silencing Complex (RISC). However, western blot-analysis of AGO2 protein levels in control cultures and cultures that expressed full-length TRIM71 or an NHL deficient form of TRIM71 protein revealed no significant changes in AGO2 expression across these conditions (Figure EV3B). In sum, TRIM71 expression in cochlear supporting cells/Kölliker’s cells does not reduce the expression of mature *let-7s*, nor does it reduce AGO2 expression, suggesting that TRIM71 increases the abundance of *let-7* targets through directly binding and stabilizing their transcripts.

Among the cohort of genes that were upregulated by *TRIM71* but not by *LIN28B* expression were BMP-responsive genes (*Bmp4, Id2, Msx1, Msx2*) (Figure 3D). During development BMP-signaling is essential for cochlear pro-sensory cell fate specification (Ohyama et al., 2010), suggesting that strengthening BMP signaling could be a key mechanism through which TRIM71 enhances supporting cell plasticity. Correlating with the ability of TRIM71 variants to enhance cell proliferation and hair cell formation, we found that expression of full-length as well as RING-domain deficient TRIM71 protein increased the expression of BMP-responsive genes (*Bmp4, Id2* and *Id3*), whereas no changes in the expression of BMP-responsive genes was observed in response to expression of NHL deficient (Figure 3G) or Coil-Coil deficient TRIM71 proteins (Figures EV3C). We next analyzed the level of phosphorylated (p)-SMAD1/5/9 proteins, an indicator of BMP signal strength, in control organoids and organoids that expressed full-length or NHL-deficient TRIM71 protein. As positive controls we also analyzed protein levels of TRIM71 target genes *Hmga2* and *Lin28b*. As anticipated, our analysis revealed that *TRIM71* expression increased BMP signaling in cochlear organoids and showed that TRIM71’s positive effect on BMP signaling requires a functional NHL domain (Figure 3EVB).

### TRIM71 promotes the de-differentiation of cochlear supporting cells/Kölliker’s cells

The observed upregulation of progenitor genes and the downregulation of supporting cell/Kölliker’s cell genes, suggests that *TRIM71* expression may reprogram supporting cells/Kölliker’s cells into progenitor-like cells. To gain further insights, we used immunostaining to analyze the changes in supporting cell and progenitor-specific proteins on a cellular level. To be able to identify differentiated/de-differentiated supporting cells/Kölliker’s cells we co-stained organoids for the Notch ligand Jagged1 (JAG1), which is selectively expressed in cochlear supporting cells, Kölliker’s cells and their precursors (Morrison et al., 1999). As progenitor-specific protein we selected the pro-sensory-specific protein HMGA2, which at stage P5 is near undetectable in cochlear supporting cells and Kölliker’s cells (Li et al., 2022). Our analysis revealed that in control organoids less than 5% of JAG1^+^ mCherry^+^ cells expressed HMGA2, while in *TRIM71*-expressing organoids more than 50% of JAG1^+^ mCherry^+^ cells expressed HMGA2, (Figures 4A, B). Conversely, in control organoids 90% of JAG1^+^ mCherry^+^ cells also expressed the supporting cell-specific protein S100A1, while in TRIM71-expressing organoids less than 30% of JAG1^+^ mCherry^+^ cells expressed S100A1 (Figures 4C, D). Potential candidates for driving TRIM71-induced changes in supporting cell-specific gene expression are the pro-differentiation transcription factors NFIB and ZBTB20 (Chen et al., 2017, Medeiros de Araújo et al., 2021). Immunostaining revealed that in the early postnatal cochlea ZBTB20 protein is selectively expressed in cochlear supporting cells, while NFIB protein is more broadly expressed in cochlear supporting cells and surrounding non-sensory epithelial cells including Kölliker’s cells (Figures EV4A, B). Furthermore, our experiments revealed that in organoid culture, *TRIM71*-expressing cells expressed NFIB and ZBTB20 at a much lower level compared to control cells and that NFIB or ZBTB20 expression was negatively correlated with hair cell-fate induction, and only cells with no or low NFIB or ZBTB20 expression co-expressed Atoh1-nGFP (Figures 4E, F) (Figures EV4C, D).

**Figure 4.**
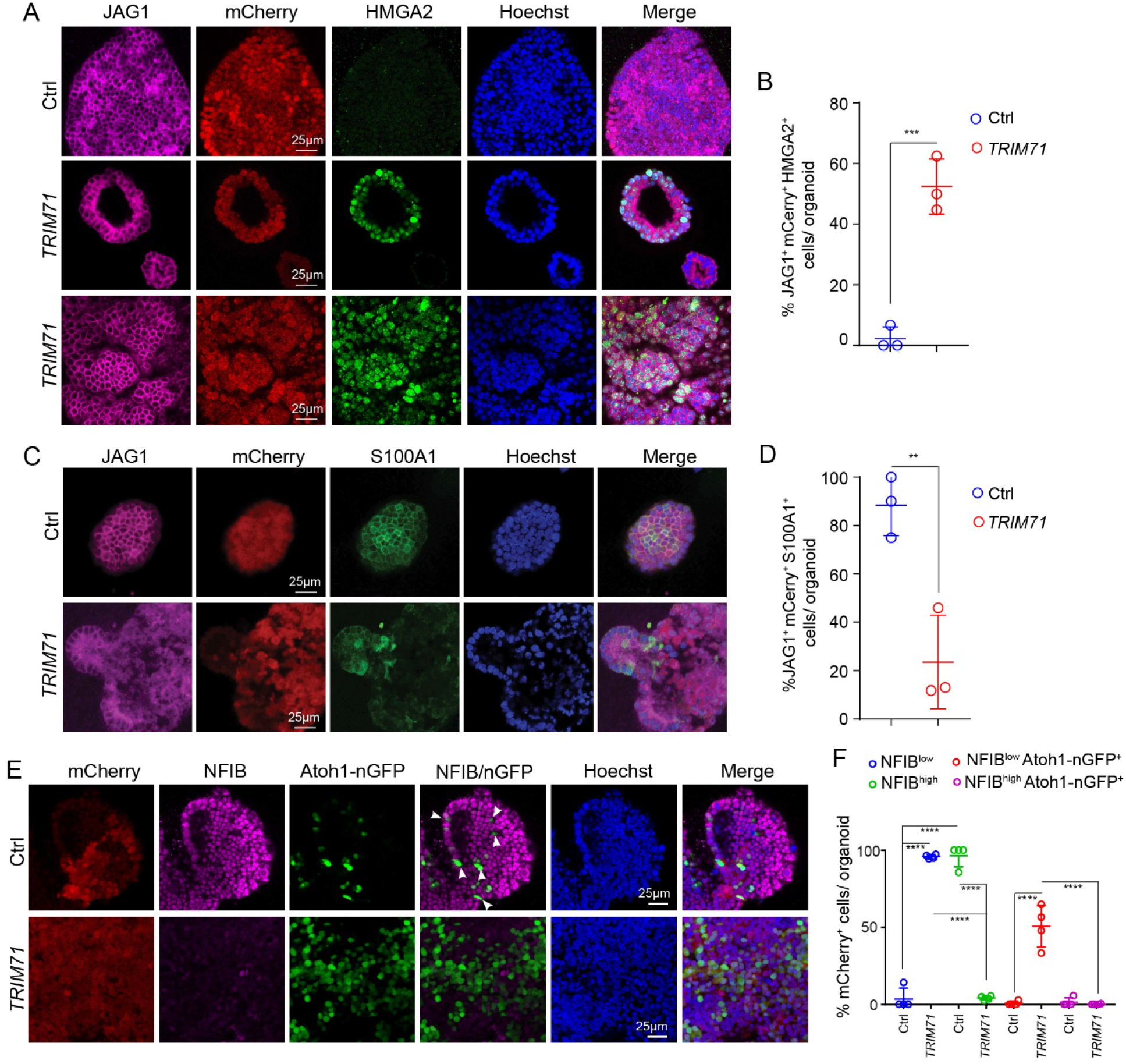
*TRIM71* promotes de-differentiation of cochlear supporting cells/Kölliker’s cells. **(A)** Confocal images showing JAG1 (magenta) and HMGA2 (green) expression in P5 control and *TRIM71*-expressing cochlear organoids at 10 days of expansion. **(B)** Quantification of JAG1^+^ mCherry^+^ HMGA2^+^ cells per organoid in (A) (n=3, two independent experiments). **(C)** Confocal images showing JAG1 (magenta) and S100A1 (green) expression in P5 control and *TRIM71-* expressing cochlear organoids at 10 days of expansion. **(D)** Quantification of JAG1^+^ mCherry^+^ S100A1^+^ cells per organoid in (A) (n=3, two independent experiments). **(E)** Confocal images showing NFIB (magenta) and Atoh1-nGFP (green) expression in P5 control and *TRIM71-* expressing cochlear organoids after 2 days in differentiation. **(F)** Quantification of NFIB^low^, NFIB^high^, NFIB^low^Atoh1-nGFP^+^, NFIB^high^ Atoh1-nGFP^+^ cells in control and *TRIM71*-expressing organoids (n=4, two independent experiments). Individual data points represent the average value per animal. One-way ANOVA with Tukey’s correction was used to calculate P values in (B) and (D). Two-way ANOVA with Tukey’s correction was used to calculate P values in (F). \**P* ≤ 0.05, \*\**P* < 0.01, \*\*\**P* < 0.001 and \*\*\*\**P* < 0.0001.

### TRIM71 and LIN28B are equally potent enhancers of supporting cell plasticity

We have previously shown that transgenic overexpression of human *LIN28B* (in the presence of FST or TGFBR inhibitor 616452) restores the hair cell-forming potential of P5 cochlear supporting cells/Kölliker’s cells (Li and Doetzlhofer, 2020, Li et al., 2022). To compare the potency of TRIM71 and LIN28B in promoting hair cell formation, we infected cochlear epithelial cells from stage P5 Atoh1-nGFP transgenic mice with lentiviral particles that expressed mCherry (control), or co-expressed mCherry with *TRIM71* or *Lin28b* or both (Figures 5A, B) (Figure EV5A). After 2 days of differentiation more than 10% of mCherry^+^ organoids in *TRIM71* or *Lin28b-*expressing cultures contained clusters of Atoh1-nGFP^+^ cells, while control cultures lacked Atoh1-GFP^+^ cell clusters. Moreover, we found that in cultures that were infected with both *TRIM71* and *Lin28b-*expressing viral particles about 25% of mCherry^+^ organoids contained Atoh1-GFP^+^ cells (Figures 5B, C). To compare the effects that the expression of *TRIM71* or *Lin28b* or both have on supporting cell reprogramming we analyzed *Hmag2, Nfib* and *Zbtb20* mRNA abundance using RT-qPCR. Our analysis revealed that individual expression of *TRIM71 or Lin28b* decreased the expression of *Zbtb20* and *Nfib* by more than 2-fold, but *Zbtb20* and *Nfib* expression was not further decreased by the co-expression of *TRIM71* and *Lin28b* (Figure 5E). Similar results were obtained analyzing the expression of *Nfib* family members *Nfia, Nfic* and *Nfix* (Figure EV5B). By contrast, co-expression of *TRIM71* and *Lin28b* further increased *Hmga2* expression by 2-fold compared to *TRIM71* or *Lin28b* alone (Figure 5D). In sum our analysis reveals that TRIM71 and LIN28B are equally potent enhancers of supporting cell reprogramming and their co-expression has a strong additive effect on hair cell formation.

**Figure 5.**
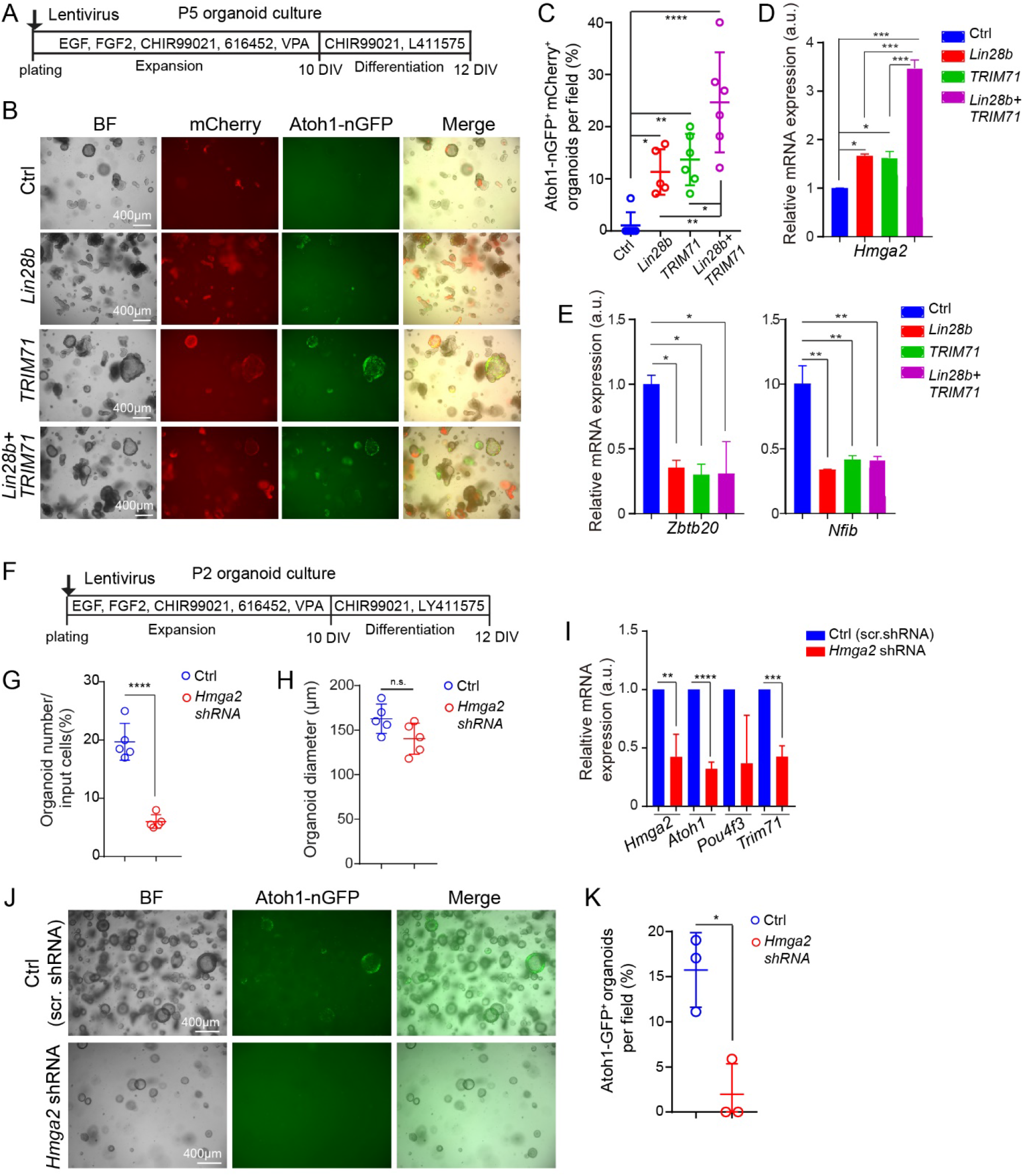
LIN28B-TRIM71 target *Hmga2* is necessary for cochlear supporting cell re-entry and subsequent hair cell formation. **(A-E)** LIN28B enhances TRIM71’s positive effect on supporting cell reprogramming and hair cell formation. **(A)** Experimental scheme. **(B)** Bright field (BF) and red and green fluorescent images of P5 cochlear organoid cultures infected with control (Ctrl) virus, or virus that expressed mouse *Lin28b*, human *TRIM71* or both (*Lin28b+TRIM71*) at 2 days of differentiation. MCherry (red) marks infected cells and Atoh1-nGFP (green) marks nascent hair cells. **(C)** Percentage of mCherry^+^Atoh1-nGFP^+^ organoids in (B). **(D-E)** RT-qPCR-analysis of progenitor (*Hmga2*) (D) and supporting cell-specific (*Zbtb20, Nfib*) (E) gene expression in control, *Lin28b, TRIM71* or *Lin28b+ TRIM71-*expressing organoids after 10 days of expansion. Individual date points represent the average value per animal. **(F-K)** Loss of *Hmga2* diminishes the mitotic and hair cell-forming potential of cochlear supporting cells/Kölliker’s cells. (**F**) Experimental scheme. (**G**) Colony forming efficiency in *Hmga2* knockdown and control cultures at 8 days in vitro (DIV). (**H**) Organoid diameter in *Hmga2* knockdown and control cultures at 8 DIV. (**I**) RT-qPCR analysis of hair cell (*Aoth1* and *Pou4f3*) and pro-sensory-specific (*Trim71*) gene expression (n=3, two independent experiments). *Hmga2* expression was analyzed to confirm knockdown. (**J**) Low power Bright field (BF) and green fluorescent images (Atoh1-nGFP) of *Hmga2* knockdown and control cultures. (**K**) Quantification of Atoh1-nGFP^+^ organoids in (J). Two-way with Tukey’s correction was used to calculate P values in (C-E) and one-way ANOVA with Tukey’s correction was used to calculate P values in (J), (H), (I) and (K). \**P* ≤ 0.05, \*\**P* < 0.01, \*\*\**P* < 0.001 and \*\*\*\**P* < 0.0001.

### LIN28B-TRIM71 target *Hmga2* is required for supporting cells/Kölliker’s cells and subsequent hair cell-formation

The TRIM71 and LIN28B-regulated gene *Hmga2* encodes a chromatin architectural protein that functions to maintain self-renewal capacity (Vignali and Marracci, 2020). To address the function of HMGA2 in hair cell regeneration we first examined whether *Hmga2 is* necessary for cell cycle re-entry, cell propagation and hair cell formation in P2 cochlear organoid culture (Figure 5F). To disrupt *Hmga2* gene function, we infected cochlear epithelial cells prior plating with lentivirus that expressed a short-hairpin RNA (shRNA) targeting endogenous *Hmga2* (Winslow et al., 2011). Control cultures were infected with lentivirus expressing a short-hairpin RNA with a scrambled sequence (scr). After 8 days of expansion, we found that *Hmga2-*knock-down cultures contained on average 4-fold fewer organoids than control cultures (Figure 5G). The average organoid size in *Hmga2*-knockdown cultures was, while mildly reduced, not significantly different from control cultures (Figure 5H). RT-qPCR confirmed the reduction in *Hmga2* expression in *Hmga2-* knockdown cultures compared to control cultures and revealed a significant reduction in the expression of *Trim71* and *Atoh1* (Figure 5I), indicating defects in reprogramming and hair cell-fate induction respectively. We next analyzed the rate of hair cell formation in *Hmga2* knockdown and control organoid cultures. We found that the percentage of Atoh1-GFP^+^ organoids in *Hmga2* knockdown cultures was significantly reduced compared to control cultures, indicating that *Hmga2* is required for hair cell formation (Figures 5J, K). We next analyzed whether overexpression of HMGA2 restores the capacity of stage P5 cochlear epithelial cells to form and grow large hair cell-containing organoids (Figure S4A). Our analysis of organoid formation efficiency and organoid growth revealed no differences between HMGA2-overexpressing cultures and control cultures (Figures S4B, C), indicating that HMGA2 overexpression is not sufficient to restore the mitotic capacity of P5 cochlear supporting cells/Kölliker’s cells. Next, we analyzed the hair cell-forming capacity of HMGA2-overexpressing and control cultures. We found that both HMGA2-overexpressing cultures and control cultures lacked Atoh1-nGFP^+^ organoids after 2 days of differentiation, indicating that HMGA2 overexpression is not sufficient to restore the hair cell regenerative capacity of P5 cochlear supporting cells/Kölliker’s cells (Figures S4D, E). In sum, our findings indicate that *Hmga2* function is required for cell cycle re-entry (organoid formation) and hair cell formation at neonatal stages but is not sufficient to restore mitotic and hair cell regenerative at later stages.

## Discussion

The loss of cochlear hair cells due to loud noise, viral infection, exposure to ototoxic substances or aging, to name only a few, is permanent, resulting in hearing deficits and deafness. Recent studies in mice have shown that immature cochlear supporting cells do have some latent capacity for hair cell regeneration (reviewed in (Zhang et al., 2020)). However, their ability to form hair cells rapidly declines as cochlear supporting cells undergo maturation. Direct reprogramming strategies such as expressing the hair cell-competence factor ATOH1 by itself (Kelly et al., 2012, Liu et al., 2012) or in combination with other hair cell-specific transcriptions factors (e.g. POU4F3, GFI1) (Walters et al., 2017) (Chen et al., 2021, Sun et al., 2021) is highly successful in converting cochlear supporting cells into hair cells at neonatal stages but thus far these strategies had no or only very limited success at later stages (Kelly et al., 2012, Liu et al., 2012, Lee et al., 2020, Sun et al., 2021).

We recently showed that rising levels of *let-7* miRNAs during cochlear maturation are a considerable barrier for hair cell regeneration and found that the mitotic and hair cell regenerative potential of cochlear supporting cells can be enhanced by overexpression of the RNA-binding protein and *let-7* antagonist LIN28B (Li and Doetzlhofer, 2020, Li et al., 2022). Here, using an organoid culture platform, we find that at embryonic stages the LIN28B/*let-7* target gene *Trim71* is essential for pro-sensory cell self-renewal and hair formation, and we show that expression of *TRIM71* at postnatal stages enhances the mitotic and hair cell regenerative potential of cochlear supporting cells (and Kölliker’s cells) to a similar extent as *Lin28b*. The strong additive effect on hair cell formation that is observed when *TRIM71* and *Lin28b* are co-expressed indicates a strong, but non-comprehensive functional overlap between them, but also non-redundant contributions, which is consistent with the findings of our RNA-seq data analysis.

TRIM71 is a potent post-transcriptional regulator, having a dual function as an ubiquitin ligase (Rybak et al., 2009) and RNA binding protein (Loedige et al., 2013). Previous studies suggested that TRIM71-mediated ubiquitination of the tumor suppressor protein p53 (Nguyen et al., 2017) and the FGF regulatory protein SHCBP1 (Chen et al., 2012) may be critical for maintaining embryonic stem cells and neural progenitor cells in an undifferentiated, proliferative state. However, our data indicates that TRIM71’s ubiquitination activity is dispensable for enhancing the mitotic and regenerative potential of cochlear supporting cells. We found that TRIM71 mutant protein that lacked E3 ubiquitin ligase activity (ΔRING) was equally capable as wild type TRIM71 in enhancing the mitotic and hair cell regenerative capacity of cochlear supporting cells. By contrast, TRIM71 mutant proteins that lack the NHL domain or the Coiled-Coil domain, which are essential for RNA-binding and protein-protein interactions failed to stimulate cochlear supporting cell proliferation. Our data is consistent with a recent study that revealed that mice homozygous for a point mutation in TRIM71’s NHL domain (R608H) have similar brain defects as *Trim71* homozygous knock out mice (Duy et al., 2022). Furthermore, the direct binding of TRIM71 to the mRNAs of the cell cycle inhibitor p21 (CDKN1A) and the pro-differentiation factor EGR1 have been shown to be essential for promoting proliferation and cell reprogramming respectively (Torres-Fernandez et al., 2019) (Worringer et al., 2014).

Recent studies revealed that TRIM71 interferes with the expression and function of mature *let-7* miRNAs through enhancing LIN28A and LIN28B-mediated repression of *let-7* biogenesis (Torres Fernandez et al., 2021) and through limiting the expression of AGO2, an essential component of miRNA-mediated gene silencing (Liu et al., 2021). Indeed, our transcriptomic profiling revealed that many of the up-regulated genes that were shared between TRIM71 and LIN28B were known or predicted targets of *let-7*-mediated gene silencing. However, our analysis revealed that TRIM71 did not reduce the expression of mature *let-7* miRNAs in cochlear organoids, nor did the presence of TRIM71 reduce AGO2 protein expression in cochlear organoids, suggesting that TRIM71 may increase the expression of *let-7* target genes by directly binding to their mRNA and stabilizing their transcript.

How does TRIM71 enhance the mitotic and regenerative potential of cochlear supporting cells? Based on our data we propose that TRIM71 reprograms cochlear supporting cells into progenitor-like cells. Supporting the idea of de-differentiation, we find that a subset of supporting cell-specific genes is rapidly downregulated in the presence of TRIM71 or LIN28B. This includes the transcription factor ZBTB20 and members of the NFI family of transcription factors (NFIA, B, C, X). ZBTB20 and NFI transcription factors function as pro-differentiation factors in glial cell types (Medeiros de Araújo et al., 2021, Chen et al., 2017) and it is tantalizing to speculate that the TRIM71 or LIN28B-induced downregulation of *Nfia,b,c,x* and *Zbtb20* expression may be a key step in priming cochlear supporting cells for hair cell-fate induction. The notion that NFI factors and ZBTB20 may interfere with hair cell fate induction is supported by our finding that high NFIB and ZBTB20 protein expression is negatively correlated with hair cell fate induction (Atoh1-GFP expression) in cochlear supporting cells. Future studies are warranted to address the role of ZBTB20 and NFI-type transcription factors in cochlear supporting cell differentiation and plasticity.

Among the genes that were upregulated by the expression of *TRIM71* or *Lin28b* or both was the *let-7* target gene *Hmga2*. HMGA2, a member of the high-mobility group HMGA family, is highly expressed in fetal and adult stem cells where it acts to maintain self-renewal potential (Copley et al., 2013, Nishino et al., 2008). In the developing cochlea *Hmga2* expression is confined to pro-sensory cells (Golden et al., 2015) and we recently found that at neonatal stages HMGA2 expression is upregulated in apical cochlear supporting cells in response to hair cell damage, suggesting that *Hmga2* re-activation may be part of an endogenous regenerative response (Li et al., 2022). Indeed our data indicates that the presence of HMGA2 is essential for the ability of cochlear supporting cells to proliferate and generate hair cells. We show that while HMGA2 is not sufficient by itself to enhance the mitotic and hair cell regenerative potential of cochlear supporting cells, loss of *Hmga2* disrupts the ability of cochlear supporting cells (and Kölliker’s cells) to re-enter the cell cycle and form hair cells. How does HMGA2 facilitate cell cycle re-entry? HMGA2 has been shown to disrupt the binding of the retinoblastoma protein (RB1) to the S-phase specific transcription factor E2F1, leading to the activation of E2F1, a step essential for cell cycle re-entry (Fedele et al., 2006). Why does HMGA2 overexpression fail to boost supporting cells plasticity? HMGA2 and other HMGA proteins do not directly regulate transcriptional activity, but rather control gene expression by changing chromatin structure and acting as co-factor for transcription factors (Vignali and Marracci, 2020). Thus, it is likely that changes in transcription factors expression/activity at the onset of cochlear maturation may limit HMGA2’s ability to enhance the regenerative potential of cochlear supporting cells.

TRIM71 and its downstream target HMGA2 are critical regulator of stem cell behavior (Mitschka et al., 2015, Zhu et al., 2011, Nishino et al., 2008), and both are known to enhance the reprogramming of adult somatic cells into induced pluripotent stem cells and neural stem cells (Worringer et al., 2014, Yu et al., 2015). However, TRIM71’s role in cell/tissue regeneration is much less well understood. In the nematode *C-elegans, Lin-41 (Trim71* orthologue) acts as a regeneration-promoting factor in young anterior ventral microtubule axons, whereas in older neurons high levels of *let-7* inhibit regeneration by down-regulating LIN-41 (Zou et al., 2013). Our work reveals that TRIM71 ability to stimulate regeneration is evolutionary conserved and thus lays the foundation for further studies investigating the role of *let-7*-TRIM71 axis in cell/tissue regeneration in mammalian and non-mammalian vertebrates. Furthermore, it will be of interest to determine whether TRIM71 facilitates cell regeneration in other sensory tissues such as the vestibular inner ear and the retina.

A limitation of our study is that we have yet to show whether TRIM71 expression would be effective in reprogramming fully mature cochlear supporting (P12 and older) into hair cell progenitor-like cells and whether reactivation of TRIM71 or co-activation of TRIM71 and LIN28B could bolster current regenerative strategies, such ATOH1 overexpression and promote cochlear hair cell formation in vivo, especially at adult stage in mammals.

## Materials and Methods

### Mouse breeding and genotyping

All experiments and procedures were approved by the Johns Hopkins University Institutional Animal Care and Use Committees protocol, and all experiments and procedures adhered to National Institutes of Health-approved standards. The *Atoh1-nGFP* transgenic (tg) (MGI:3703598) mice were provided by Jane Johnson, University of Texas Southwestern Medical Center, Dallas, TX (Lumpkin et al., 2003). The *Col1a-TRE-LIN28B* (MGI:5294612) mice were provided by George Q. Daley, Children’s Hospital, Boston, MA (Zhu et al., 2011). *Trim71^f/f^* mice were provided by Waldemar Kolanus (LIMES Institute, Universität Bonn, Germany). *TetO-cre* (No.006234) and *R26^rtTA*M2^* (No. 006965) mice were purchased from Jackson Laboratories (Bar Harbor, ME). We crossed *TetO-cre tg; Tnm71^f/f^* males/females with *Trim71^f/f^; R26^rtTA*M2/rtTA*M2^* males/females to obtain *Trim71 KO (TetO-cre tg; R26r^tTA*M2/+^; Trim71^f/f^*) and control littermates (*R26^rtTA*M2/+^; Trim71 ^f/f^*). To induce Cre expression, doxycycline (dox) was delivered to time-mated females via ad libitum access to feed containing 2 g/kg dox. Mice were genotyped by PCR as previously published. Genotyping primers are listed in table S4. Mice of both sexes were used in this study. Embryonic development was considered as E0.5 on the day a mating plug was observed.

### Lentiviral vectors and production

Expression plasmids for full-length and mutant TRIM71 (ΔRING, Δcoiled-coil, and ΔNHL) proteins were generated by sub cloning HA-tagged human *TRIM71* and its mutant variants sequences from *pMXS-hs-3xHA-TRIM71* (*Addgene, no. 52717*), *pMXs-hs-3xHA-deltaRING-TRIM71* (Addgene, no. 52718), *pMXs-hs-3xHA-delta Coiled Coil-TRIM71* (Addgene, no. 52720) and *pMXs-hs-3xHA-delta6xNHL-TRIM71* (Addgene, no. 52722) into *pCDH-EF1a-eFFly-T2A-mCherry* (Addgene, no. 104833) replacing eFFly with restriction enzymes XbaI (NEB, no. R0145) and EagI-HF (NEB, no. R3505). TRIM71-R608H expression plasmid was generated by cloning DNA fragment-R608H (Integrated DNA Technology) (table S5) into *pCDH-EF1a-3xHA-TRIM71-T2A-mCherry* plasmid using SmaI (NEB, no. R0141) and EcoRI-HF (NEB, no. R3101). TRIM71-R796H expression plasmid was generated by cloning DNA fragment-R796H (Integrated DNA Technology) (table S5) into *pCDH-EF1a-3xHA-TRIM71-T2A-mCherry* using EcoRI-HF and EagI-HF. TRIM71-R608H+R796H expression plasmid was generated by cloning DNA fragment-R608H into *pCDH-EF1a-3xHA-TRIM71-(R796H)-T2A-mCherry* using SmaI and EcoRI-HF. Lentiviral control construct was obtained by replacing the eFFly in *pCDH-EF1a-eFFly-T2A-mCherry* with a 3xHA sequence using restriction enzymes XbaI and BamHI-HF (NEB, no. R3136). The *pFUWFLAG-Lin28b-F2A-mCherry* construct expressing mouse *Lin28b* under the control of human ubiquitin C promoter was previously described (Golden et al., 2015). Lentiviral constructs for HMGA2 overexpression were obtained by sub cloning human *HMGA2* from *pMXS-hs-HMGA2* (Addgene, no. 52727) into *pCDH-EF1a-eFFly-mCherry* replacing eFFly using restriction enzymes XbaI and BamHI-HF. Lentiviral *shHmga2* construct was purchased from Addgene (no. 32399). Lentiviral packaging was conducted as previously described (Li et al., 2022).

### Organoid culture

Cochlear organoid cultures were conducted as previously described (Li and Doetzlhofer, 2020). Briefly, cochlear epithelia were enzymatically isolated from stage P2 or P5 mice, reduced to single cells, and for lentiviral infection, cells were mixed and centrifuged with lentiviral particles at 600 g for 30 minutes. Otherwise, cells were immediately resuspended in expansion medium [DMEM/F12 (Corning, no. 10–092-CV), N-2 supplement (1x, ThermoFisher, no. 17502048), B-27 supplement (1x, ThermoFisher, no. 12587010), EGF (50 ng/mL, Sigma-Aldrich, no. SRP3196), FGF2 (50 ng/mL, ThermoFisher, no. PHG0264), CHIR99021 (3 μM, Sigma-Aldrich, no. SML1046), VPA (1 mM, Sigma-Aldrich, no. P4543), 616452 (2 μM, Sigma-Aldrich, no. 446859-33-2) and penicillin (100 U/mL, Sigma-Aldrich, no. P3032)] mixed 1:1 with Matrigel (Corning, no. 356231), and cultured in expansion media for 10 days. To induce hair cell formation organoids were cultured in differentiation medium [DMEM/F12, N2 (1x), B27 (1x), CHIR99021 (3 μM) and LY411575 (5 μM, Sigma-Aldrich, no. SML0506)].

### RNA extraction, RT-qPCR and TaqMan assay

Organoids were harvested using Cell Recovery Solution. Total RNA from organoids/tissue was extracted using the miRNeasy Micro Kit (QIAGEN). MRNA was reverse transcribed into cDNA using the iScript cDNA synthesis kit (Bio-Rad). Q-PCR was performed on a CFX-Connect Real Time PCR Detection System using SYBR Green Master Mix reagent (Applied Biosystems). Gene-specific primers used are listed in table S6. *Rpl19* was used as an endogenous reference gene. Relative gene expression was calculated using ΔΔCT method. To quantify the expression of mature *let-7a-5p, let-7d-5p, let-7g-5p, let-7i-5p* transcripts, predesigned TaqMan Assays (Applied Biosystems) were used according to manufacturer’s instructions. The snoRNA U6 was used as an endogenous reference gene for TaqMan-based miRNA measurements.

### RNA sequencing and data analysis

Cochlear epithelial cells from stage P5 wild-type mice were used to establish organoid culture. Before plating, cells were infected with control lentivirus (3xHA-T2A-mCherry) or lentivirus expressing human *TRIM71* (3xHA-TRIM71-T2A-mCherry). For each condition, three independent cultures were established. At 10 days of expansion, RNA was extracted, and samples were processed using Illumina’s TruSeq stranded Total RNA kit, per manufacturer’s recommendations, using the UDI indexes. The samples were sequenced on the NovaSeq 6000, paired end, 2×50 base pair reads. Kallisto (v0.46.1) was used to pseudo-align reads to the reference mouse transcriptome and to quantify transcript abundance. The transcriptome index was built using the Ensembl Mus musculus v96 transcriptome. The companion analysis tool sleuth was used to identify differentially expressed genes (DEGs). We performed a Wald test to produce a list of significant DEGs between *TRIM71* expressing sample and the control. These lists were then represented graphically using sleuth along with pheatmap and ggplot2 packages in R v1.3.1093. Gene identifier conversion, gene annotation, and enrichment analysis was conducted using Metascape (Zhou et al., 2019).

### Immunohistochemistry

Organoids were fixed in 4% paraformaldehyde for 30 minutes, permeabilized and blocked with 0.25% Triton X-100/ 10% fetal bovine serum for 30 minutes, and immuno-stained as previously described (Li and Doetzlhofer, 2020) (Li et al., 2022). Antibodies are listed in table S6.

### Cell proliferation

EdU (Thermo Fisher Scientific, no. C10338) was added to culture medium at a final concentration of 3 μM. EdU incorporation was analyzed using the Click-iTPlus EdU Cell Proliferation Kit (Thermo Fisher Scientific, no. C10637 and C10638) following the manufacturer’s recommendations.

### Cell counts

High-power confocal single-plane and z-stack images of fluorescently immuno-labeled organoids and explants were taken with 40x objective using LSM 700 confocal microscope (Zeiss Microscopy). To establish the percentage of cells within an organoid that expressed an epitope of interest, at least three images were analyzed and the average value per animal was reported. At a minimum, two independent experiments were conducted in which at a minimum three organoid cultures per genotype and treatment were established and analyzed.

### Protein lysis co-Immuno-precipitation and immunoblotting

HEK 293T cells were transiently transfected with a total of 2.5 μg of plasmid DNA. FLAG-Lin28b (Golden et al., 2015) or FLAG-LIN28A (Addgene no. 51371) constructs were co-transfected with 3xHA-TRIM71 expressing constructs using Polyethylenimine, Linear, MW 25000, transfection reagent (Polysciences, no. 23966). After 48 hours, cells were lysed in NP-40 lysis buffer (Thermo Scientific, no. J60766.AP) supplemented with protease inhibitor (Roche, no.11836170001). 1% of cell lysate was saved as input, the remaining lysate precleared for 1 hour with Protein-A Dynabeads (Thermo Scientific, no. 10001D). FLAG-M2 antibody Protein-A Dynabead conjugate was incubated with the precleared lysate overnight. Proteins bound to Protein-A Dynabeads were pulled down using a magnetic separation rack (Cell Signaling, no. 14654). After 5 washes in NP-40 lysis buffer, beads were resuspended in 4x Laemmli protein sample buffer (Bio-Rad, no.1610747). Protein separation and immunoblots were conducted as previously described (Li and Doetzlhofer, 2020). The resulting chemiluminescence was captured using x-ray films or digitally using LI COR Odyssey imaging system. The antibodies used are listed in table S7. Cochlear organoids were lysed with RIPA buffer (Sigma-Aldrich, no. R0278) supplemented with protease inhibitor (Sigma-Aldrich, no. 11697498001), phosphatase Inhibitor cocktail 2 (Sigma-Aldrich, no. P5726) and phosphatase inhibitor cocktail 3 (Sigma-Aldrich, no. P0044).

### Quantification of organoid forming efficiency, organoid diameter, and GFP^+^ cells

Low-power bright-field and fluorescent images of organoid cultures were captured with an Axiovert 200 microscope using 5x objectives (Carl Zeiss Microscopy). The organoid formation efficiency and the diameter of organoids were measured as previously described (Li and Doetzlhofer, 2020) (Li et al., 2022), and the average value per animal was reported. To establish the percentage of GFP^+^ organoids per culture, the total number of organoids and the number of GFP^+^ or GFP^+^ mCherry^+^ organoids were established by counting manually. For each genotype and treatment, three independent organoid cultures from three different animals were established and analyzed. At a minimum, two independent experiments were conducted and analyzed.

### Statistical analysis

All results were confirmed by at least two independent experiments. The sample size (n) represents the number of animals analyzed per group. Animals (biological replicates) were allocated into control or experimental groups based on genotype and/or type of treatment. To avoid bias, masking was used during data analysis. Data was analyzed using Graphpad Prism 8.0. Relevant information for each experiment including sample size, statistical tests and reported p-values are found in the legend corresponding to each figure. *P ≤ 0.05, **P < 0.01, ***P < 0.001 and ****P <0.0001. In all cases p-values ≤ 0.05 were considered significant and error bars represent standard deviation (SD).

## Acknowledgement

We thank the members of the Doetzlhofer Laboratory for the help and advice provided throughout the course of this study.

## Author’s contributions

Conceptualization: A.D., X.-J.L.

Methodology: A.D., X.-J.L, C.M., P.T.N.P, W.K.

Investigation: X.-J.L., C.M. P.T.N.P

Supervision: A.D.

Writing—original draft: A.D., X.-J.L.

Writing—review & editing: A.D., X.-J.L., C.M, P.T.N.P, W.K

## Data and materials availability

RNA sequencing data have been deposited in the Gene Expression Omnibus data repository under accession number GSE210383.

**Tables S1-S3 are provided as separate files in spreadsheet format (excel file)**

**Table S1** Wald test Ctrl vs *TRIM71* expressing

**Table S2** GO analysis Ctrl vs *TRIM71* expressing

**Table S3** Comparison *TRIM71* and *LIN28B+FST* regulated genes

## Expanded View Figures

**Figure EV2.**
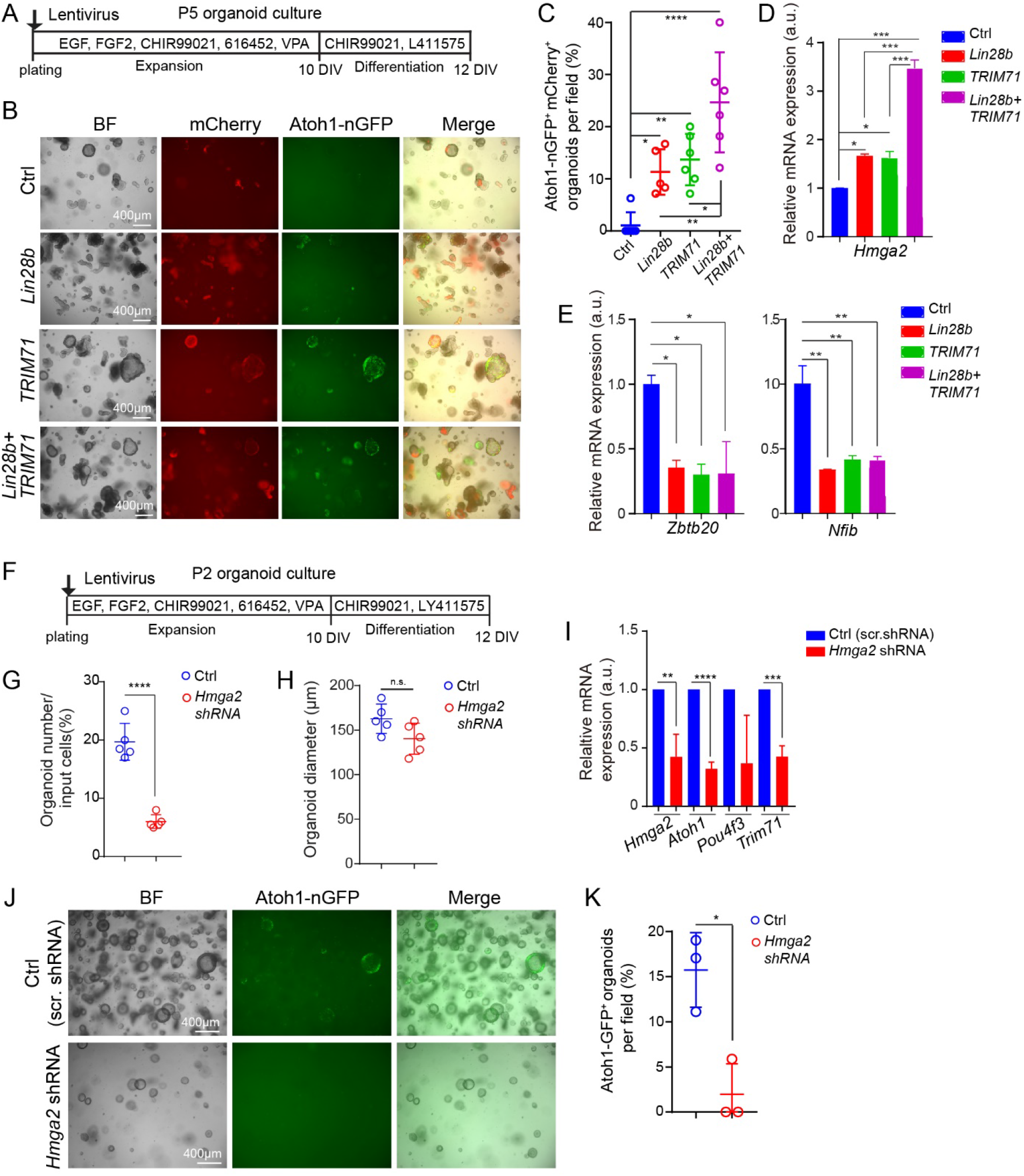
*Trim71* deficient cochlear progenitor cells have diminished capacity for self-renewal and hair cell formation. **(A)** Schematic of experimental strategy. Organoid cultures were established with cochlear epithelial cells from stage E13.5 *Trim71* KO (*TetO-Cre; R26rtTA*M2; Trim71f/f; Atoh1-nGFP*) mice and littermates that lacked *TetO-Cre* transgene (WT). Timed pregnant dam received doxycycline (dox) containing feed starting at E5.5 until tissue harvest. The *Atoh1-nGFP* expression marks nascent hair cells. **(B)** Bright field (BF) images of E13.5 wild type and *Trim71* KO organoids at 7 and 10 days of expansion. **(C)** Organoid diameter in (B) (n=3, two independent experiments). **(D)** Colony forming efficiency in (B) (n=3, two independent experiments). **(E)** Cell proliferation in E13.5 wild type and *Trim71* KO organoids. An EdU pulse was given at 7 days of expansion and EdU incorporation (red) was analyzed 1.5 hours later. Hoechst labels cell nuclei (blue). **(F)** Percentage of EdU^+^ cells in E13.5 wild type (blue) and *Trim71* KO (red) organoids (n = 9, three independent experiments). **(G)** Low power bright field (BF) and green fluorescent (Atoh1-nGFP) images of E13.5 wild type and *Trim71* KO organoid cultures. **(H)** Quantification of Atoh1-nGFP^+^ organoids in (G) (n=3, two independent experiments). Individual date points represent the average value per animal. Two-tailed, unpaired *t* test was used to calculate *P* values in (C), (D), (F) and (H). \*\**P* < 0.01 and \*\*\**P* < 0.001.

**Figure EV3.**
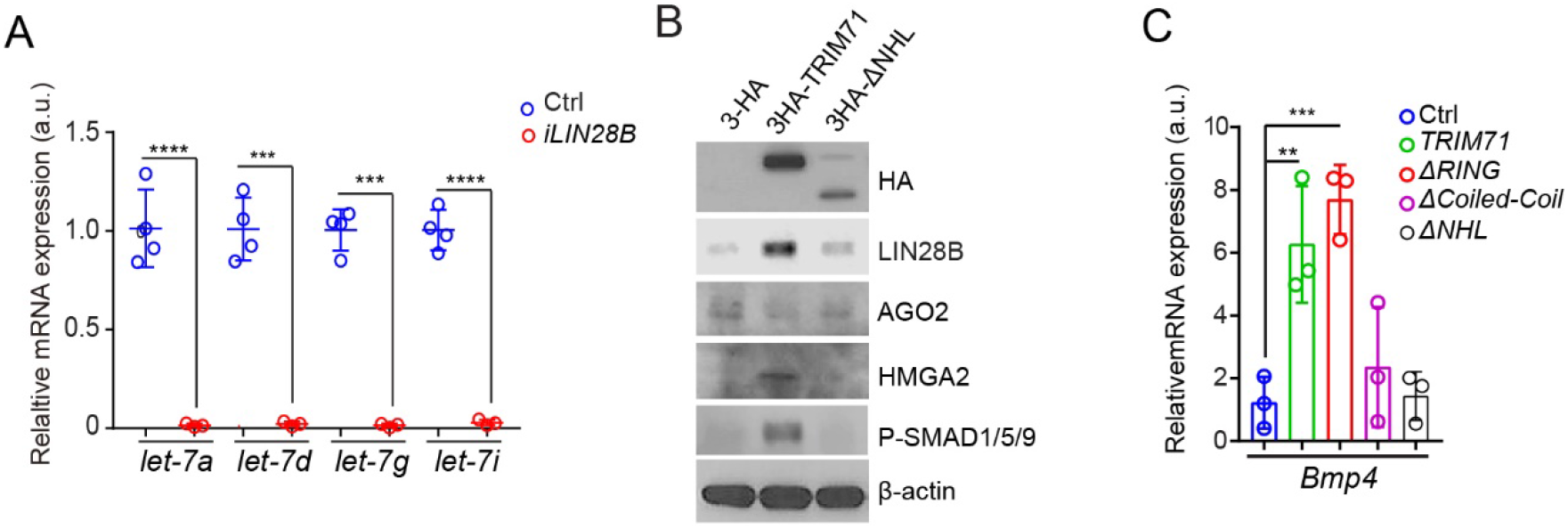
Effects of LIN28B and TRIM71 on BMP and let-7 target genes. **(A)** TaqMan assay of mature *let-7a-5p, let-7d-5p, let-7g-5p, let-7i-5p* transcripts in P5 control (Ctrl) and *LIN28B*-expressing cochlear organoids (n=3, two independent experiments). Two-tailed, unpaired *t* test was used to calculate *P* values. **(B)** Immunoblots were used to analyze endogenous LIN28B, AGO2, HMGA2, P-SMAD1/5/9 and β-actin protein levels in P5 control, TRIM71 and ΔNHL expressing cochlear organoids at 10 days of expansion. **(C)** RT-qPCR of *Bmp4* mRNA expression (n=3, two independent experiments) in P5 control (Ctrl) cochlear organoids and cochlear organoids that expressed full length (*TRIM71*), RING deficient (Δ*RING*), Coiled-Coil deficient (*ΔCoiled-Coil*) or NHL deficient (*ΔNHL*) TRIM71 protein at 10 days of expansion. Individual date points represent the average value per animal. One-way ANOVA with Tukey’s correction was used to calculate *P* values. \*\**P* < 0.01, \*\*\**P* < 0.001 and \*\*\*\**P* < 0.0001.

**Figure EV4.**
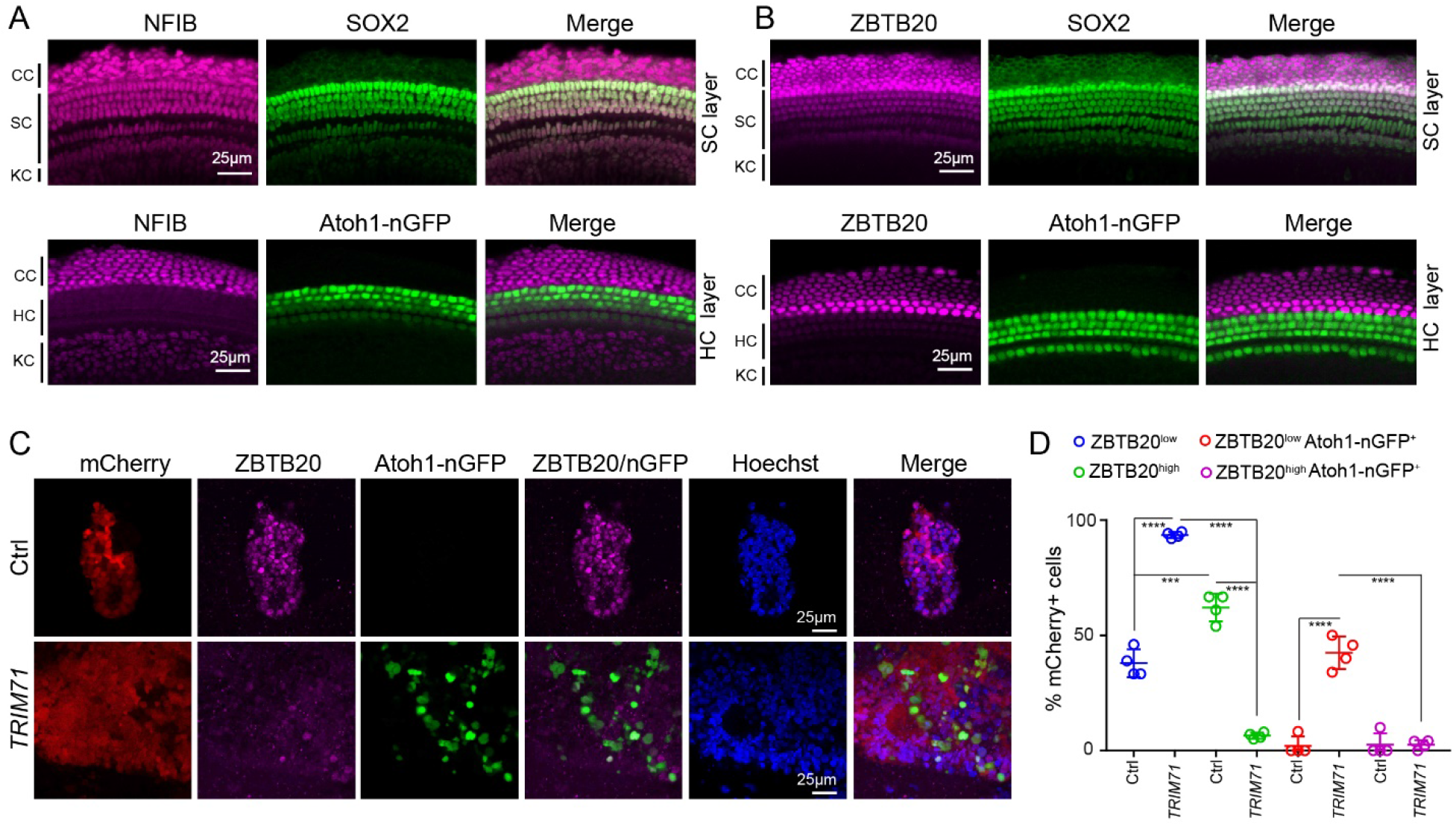
TRIM71 represses the expression of supporting cell-specific transcription factor ZBTB20. **(A-B)** Cochlear ZBTB20 and NFIB protein expression in vivo. Confocal images of the hair cell layer (HC) and supporting cell layer (SC) of cochlear sensory of stage P5 mice. SOX2 (green) marks supporting cells (SC) and Kölliker’s cells (KC) but not Claudius cells (CC). Atoh1-nGFP (green) marks hair cells (HC). **(A)** NFIB (magenta) is expressed in Claudius cells, supporting cells and Kölliker’s cells but not in hair cells. **(B)** ZBTB20 (magenta) is expressed in Claudius cells and supporting cells, but not in Kölliker’s cells or hair cells. **(C)** Confocal images of ZBTB20 protein (magenta) and Atoh1-nGFP transgene (green) expression in P5 control and TRIM71 expressing cochlear organoids after 2 days of differentiation. **(D)** Quantification of ZBTB20^low^, ZBTB20^hi^g^h^, ZBTB20^low^ Atoh1-nGFP^+^, ZBTB20^hi^g^h^ Atoh1-nGFP^+^ cells in (C) (n=4, two independent experiments). Individual date points represent the average value per animal. Two-way ANOVA with Tukey’s correction was used to calculate *P* values in D. *\*P* ≤ 0.05, *\*\*P* < 0.01, *\*\*\*P* < 0.001 and \*\*\*\**P* < 0.0001.

**Figure EV5.**
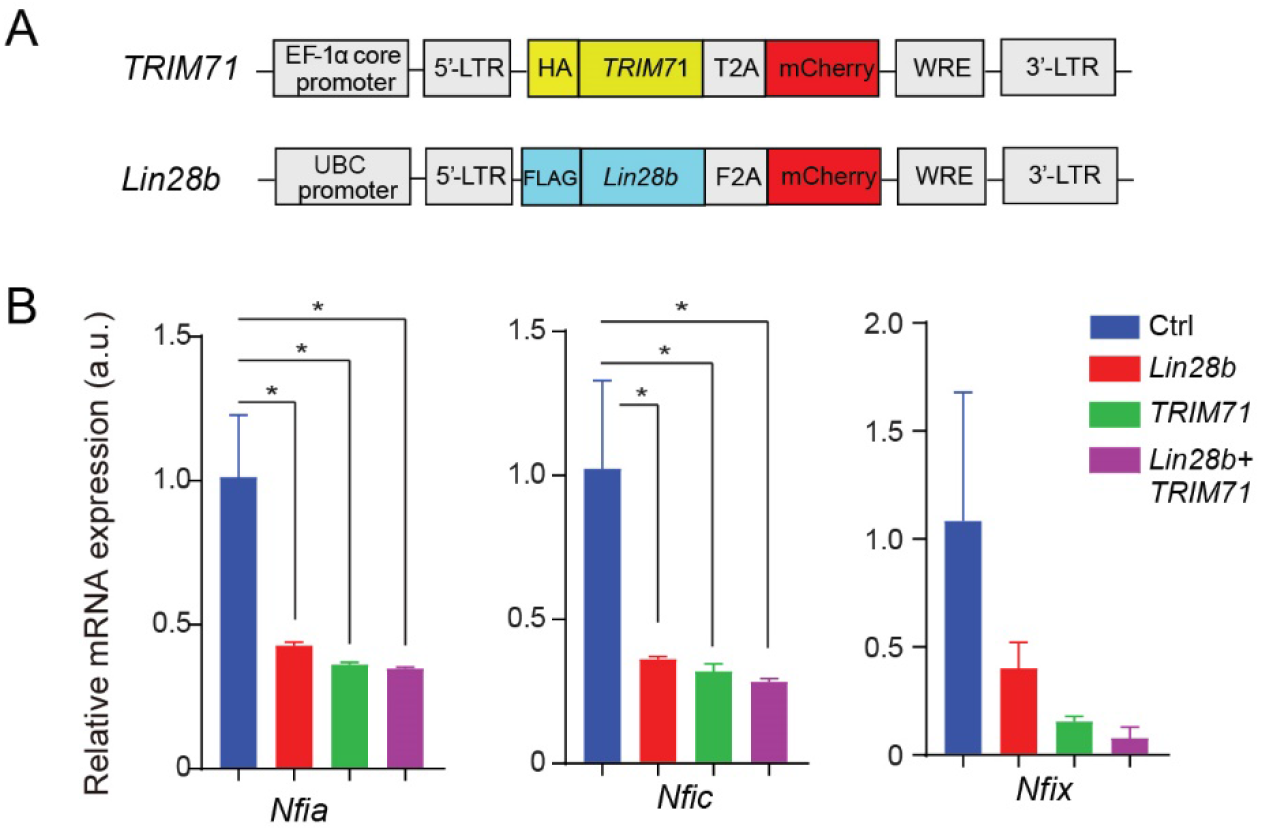
**(A)** Schematic of lentiviral expression cassettes used to express human *TRIM71* and mouse *Lin28b* in P5 cochlear organoid cultures. **(B)** RT-qPCR-analysis of *Nfia, Nfic* and *Nfix* expression in control (Ctrl), *Lin28b, TRIM71* or *Lin28b+TRIM71-expressing* organoids after 10 days of expansion. Two-way ANOVA with Tukey’s correction was used to calculate P values. \**P* ≤ 0.05,

