## Supplemental Methods and Materials for "Reactivation of the progenitor gene Trim71 enhances the mitotic and hair cell-forming potential of cochlear supporting cells"

### Supplemental Material

| Mouse line | Genotyping primers | Product size |
|---|---|---|
| R26 $^{rtTA*M2}$ | MTR: GCG AAG AGT TTG TCC TCA ACC<br>F: AAA GTC GCT CTG AGT TGT TAT<br>WTR: GGA GCG GGA GAA ATG GAT ATG | WT=650bp<br>MT=340bp |
| *Trim71$^{fl/fl}$* | LIN41-F-WT: GAA AGG AGG CTA GCC AAA GG<br>LIN41-R-KO: ATG CTG TAC GGT AGG AGT CTT CC | FL=350bp<br>WT=250bp |
| *Col1a-TRE-LIN28B* | ColA: GCA CAG CAT TGC GGA CAT GC<br>ColB: CCC TCC ATG TGT GAC CAA GG<br>ColC: GCA GAA GCG CGG CCG TCT GG | WT=300bp<br>TG=450bp |
| *Atoh1-nGFP* | EGFP1: CGA AGG CTA CGT CCA GGA GCG CAC<br>EGFP2: GCA CGG GGC CGT CGC CGA TGG GGG TGT | TG= 300bp |
| *TetO-cre* | F: GCCTGCATTACCGGTCGATGCAACGA<br>R: GTGGCAGATGGCGCGGCAACACCATT | TG:700bp |

**Table S4.** List of genotyping primers.

| | |
|---|---|
| DNA-fragment-R608H | CCTGAGCTTCGGCAGTGAGGGTGACAGCGATGGCAAGCTCTGCCACCCTT<br>GGGGTGTGAGTGTAGACAAGGAGGGCTACATCATTGTCGCCGACCGCAG<br>CAACAACCGCATCCAGGTGTTCAAGCCCTGCGGCGCCTTCCACCACAAAT<br>TCGGCACCCTGGGCTCCCGGCCTGGGCAGTTCGACCGACCAGCCGGCGT<br>GGCCTGTGACGCCTCACGCAGGATCGTGGTGGCTGACAAGGACAATCATC<br>GCATCCAGATCTTCACGTTCGAGGGCCAGTTCCTCCTCAAGTTTGGTGAG<br>AAAGGAACCAAGAATGGGCAGTTCAACTACCCTTGGGATGTGGCGGT |
| DNA-fragment-R796H | TGAGGGCAAGATCCTGGTCTCAGACACGAGGAACCACCGGATCCAGCTGT<br>TTGGGCCTGATGGTGTCTTCCTAAACAAGTATGGCTTCGAGGGGGCTCTC<br>TGGAAGCACTTTGACTCCCCACGGGGTGTGGCCTTCAACCATGAGGGCCA<br>CTTGGTGGTCACTGACTTCAACAACCACCGGCTCCTGGTTATTCACCCCGA<br>CTGCCAGTCGGCACGCTTTCTGGGCTCGGAGGGCACAGGCAATGGGCAG<br>TTCCTGCACCCACAAGGGGTAGCTGTGGACCAGGAAGGGCGCATCATTGT<br>GGCGGATTCCAGGAACCATCGGGTACAGATGTTTGAATCCAACGGCAGCT<br>TCCTGTGCAAGTTTGGTGCTCAAGGCAGCGGCTTTGGGCAGATGGACCGC<br>CCTTCCGGCATCGCCATCACCCCCGACGGAATGATCGTTGTGGTGGACTT<br>TGGCAACAATCGAATCCTCGTCTTCG |

**Table S5.** DNA fragments for cloning

| Gene | Forward Primer | Reveres Primer |
|---|---|---|
| *Ano1* | TTC CCT CTG GCT CCA CTC TTC | GGC ATC CAG GCG GAT CT |
| *Atoh1* | ATG CAC GGG CTG AAC CA | TCG TTG TTG AAG GAC GGG ATA |
| *Bmp4* | ACG TAG TCC CAA GCA TCA CC ACT | AGG GTC TGC ACA ATG GC |
| *Ccnd2* | GGG ATC CCT GTA CAC TCG AA | TTG CAG GTA CGC ACA CTC TC |
| *Cybrd1* | AGA CTG CCA TGG ACC TGG AA | CCG GCA TGG ATG GAT TTC |

| | | |
|---|---|---|
| *Fat3* | CAC AGC CCT TGA ATA CAG TGA | TGC CTT TGC ATC TCC TTC CT |
| *Fst* | GAA AAC CTA CCG CAA CGA ATG | TCC GGC TGC TCT TTG CAT |
| *Gfi1* | AGGAACGCAGCTTTGACTGT | TGAGATCCACCTTCCTCTGG |
| *Hmga2* | CAG AAG AAA GCA GAG ACC ATT GG | TTG TTG TGG CCA TTT CCT AGG T |
| *Id1* | GAA CGT CCT GCT CTA CGA CAT G | TGG GCA CCA GCT CCT TGA |
| *Id2* | AAG GTG ACC AAG ATG GAA ATC CT | CGA TCT GCA GGT CCA AGA TGT |
| *Id3* | GAG CTC ACT CCG GAA CTT GTG | CGG GTC AGT GGC AAA AGC |
| *Lgr5* | CCC CAA TGC GTT TTC TAC GT | GAA GGA CGA CAG GAG ATT GGA T |
| *Myo7a* | CCC CCT CTG AGA AGT TCG TTA A | TGT GTC CGA GTT CCG TTG AC |
| *Nfia* | GAG TCC AGG AGC AAT GAG G | CCA TTT CAT CCT CCA CAG AC |
| *Nfib* | GTG TTC AGC CAC ACC ACA TC | GAG GAT TCT TGG CAG GAT CA |
| *Nfic* | CCG GCA TGA GAA GGA CTC TAC | TTC TTC ACC GGG GAT GAG ATG |
| *Nfix* | AGG CTG ACA AGG TGT GGC | CAC TGG GGC GAC TTG TAG AG |
| *Pou4f3* | GCA CCA TCT GCA GGT TCG A | CCG GCT TGA GAG CGA TCA T |
| *Rpl19* | GGT CTG GTT GGA TCC CAA | TGC CCG GGA ATG GAC AGT CA |
| *S100a1* | TGG ATG TCC AGA AGG ATG CA | CCG TTT TCA TCC AGT TCC TTC A |
| *Sox2* | CCA GCG CAT GGA CAG CTA | GCT GCT CCT GCA TCA TGC T |
| *Trim71* | ATC GGG AGT GTG AGC TGT TG | GGC GTG AAC ATA ATG CGG TC |
| *Zbtb20* | GGC ATC TGA GGA GAA TGA GA | GTT GTG AAG GTT GAT GCT GTG |

**Table S6.** List of qPCR primers.

| **Immunostaining** | | | | |
|---|---|---|---|---|
| Reagent type | Designation | Source | Identifiers | Additional information |
| antibody | myosinVIIa rabbit polyclonal | Proteus Biosciences | Cat.# 25-6790 | 1:500 dilution |
| antibody | SOX2 goat polyclonal | Santa Cruz | Cat.# sc-17320 | 1:500 dilution |
| antibody | JAG1 goat polyclonal | Santa Cruz | Cat.# sc-6011 | 1:500 dilution |
| antibody | HMGA2 rabbit monoclonal | Cell Signaling | Cat.# 8179 | 1:500 dilution |
| antibody | S100-alpha rabbit polyclonal | Abcam | Cat.# ab11428 | 1:500 dilution |
| antibody | ZBTB20 rabbit polyclonal | Sigma | Cat.# HPA016815 | 1:500 dilution |
| antibody | NFIB rabbit polyclonal | Sigma | Cat.# HPA003956 | 1:500 dilution |
| antibody | donkey anti-rabbit IgG (H+L) Alexa Fluor 647 | ThermoFisher | Cat.# A-31573 | 1:1000 dilution |
| antibody | donkey anti-rabbit IgG (H+L) Alexa Fluor 488 | ThermoFisher | Cat.# A32790 | 1:1000 dilution |
| antibody | donkey anti-goat IgG (H+L) Alexa Fluor 647 | ThermoFisher | Cat.# A32849 | 1:1000 dilution |
| antibody | donkey anti-goat IgG (H+L) Alexa Fluor 488 | ThermoFisher | Cat.# A-11055 | 1:1000 dilution |

| | | | | |
|---|---|---|---|---|
| antibody | Biotinylated donkey anti goat | Jackson Immuno Research Lab | Cat.#705-065-147 | 1:200 dilution |
| dye | Streptavidin, Alexa Fluor 405 | Life Technologies | Cat. #S32351 | 1:200 dilution |
| stain | Hoechst 33258 solution | Sigma-Aldrich | Cat.# 94403 | 1:3000 dilution |
| **Immunoblotting** | | | | |
| Reagent type | Designation | Source | Identifiers | Additional information |
| antibody | FLAG (M2) mouse monoclonal | Sigma | Cat. # F1804 | 1:1000 dilution |
| antibody | LIN28B (mouse preferred) rabbit polyclonal | Cell Signaling | Cat.# 5422 | 1:1000 dilution |
| antibody | HMGA2 rabbit monoclonal | Cell Signaling | Cat.# 8179 | 1:1000 dilution |
| antibody | AGO2 rabbit monoclonal | Cell Signaling | Cat.# 2897 | 1:1000 dilution |
| antibody | P-SMAD1/5/9 (D5B10) rabbit monoclonal | Cell Signaling | Cat.# 13820 | 1:1000 dilution |
| antibody | β-actin mouse monoclonal | Santa Cruz | Cat.# 47778 | 1:500 dilution |
| antibody | HRP-conjugated donkey anti-mouse IgG | Jackson Immuno Research | Cat. #715-035-150 | 1:10000 dilution |
| antibody | HA (C29F4) rabbit monoclonal | Cell Signaling | Cat.# 3724 | 1:1000 dilution |
| antibody | HRP-conjugated goat anti-rabbit IgG, | Jackson Immuno Research | Cat.# 111-035-003 | 1:10000 dilution |

**Table S6.** List of antibodies, dyes and stains

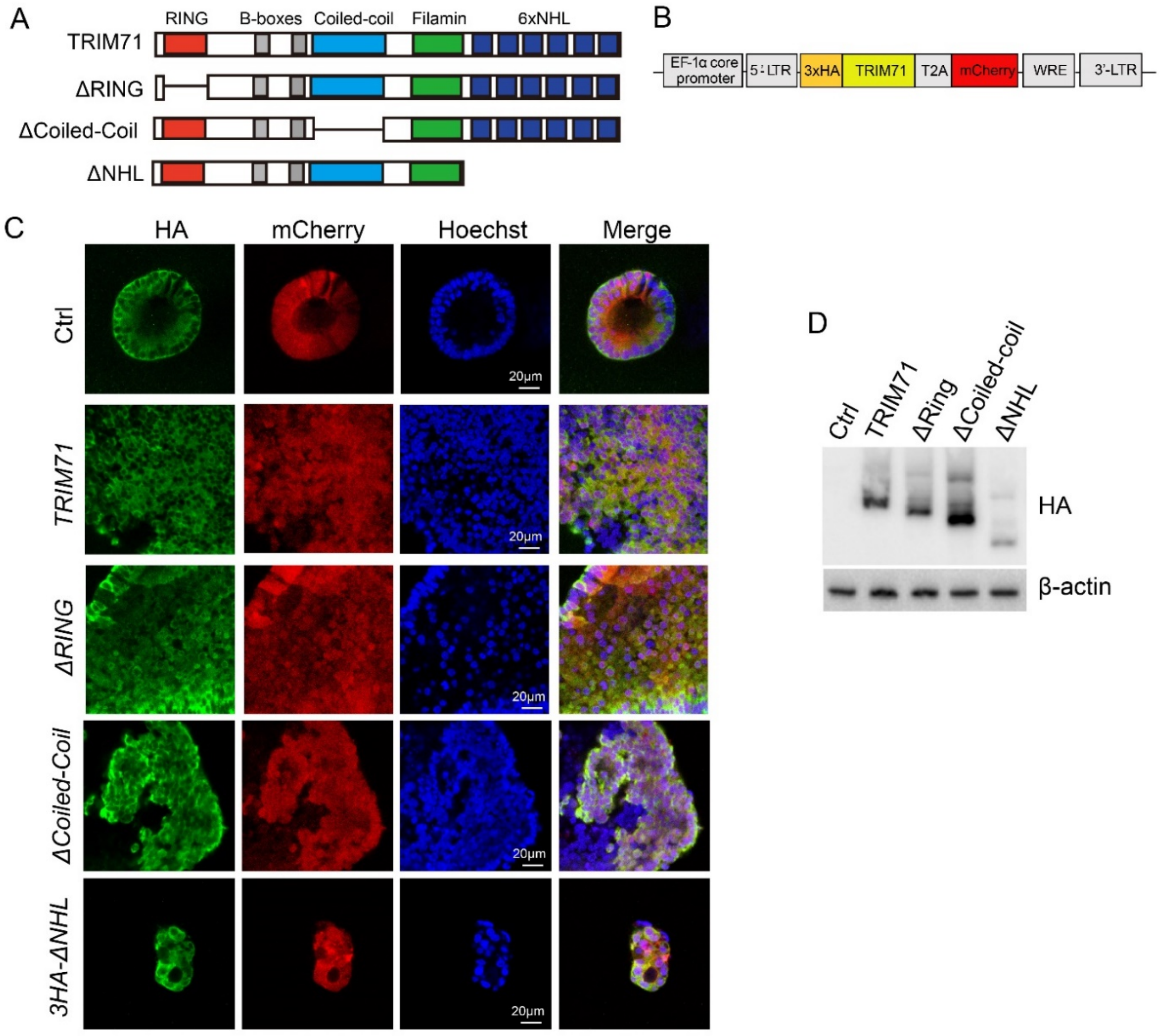


**Supplemental Figure 1. Structure of TRIM71 mutants. (A)** Schematic of human TRIM71 protein and mutant variants that lack the RING (red, ∆ RING), Coiled coil (light blue, ∆ Coiled coil) or NHL domains (dark blue, ∆ NHL). **(B)** Schematic of expression cassette. **(C)** Confocal images of P5 cochlear organoids infected with lentivirus expressing TRIM71 protein and TRIM71 mutant variants. MCherry (red) labels infected cells and HA immunostaining (green) visualizes TRIM71 protein expression. Note control cultures were infected with virus that expresses HA only. **(D)** Protein expression levels of full-length TRIM71 (TRIM71) and TRIM71 mutant variants (△RING, △Coiled-Coil and △NHL) in P5 cochlear organoids at 10 days of expansion. Shown are immunoblots against HA and β-actin.

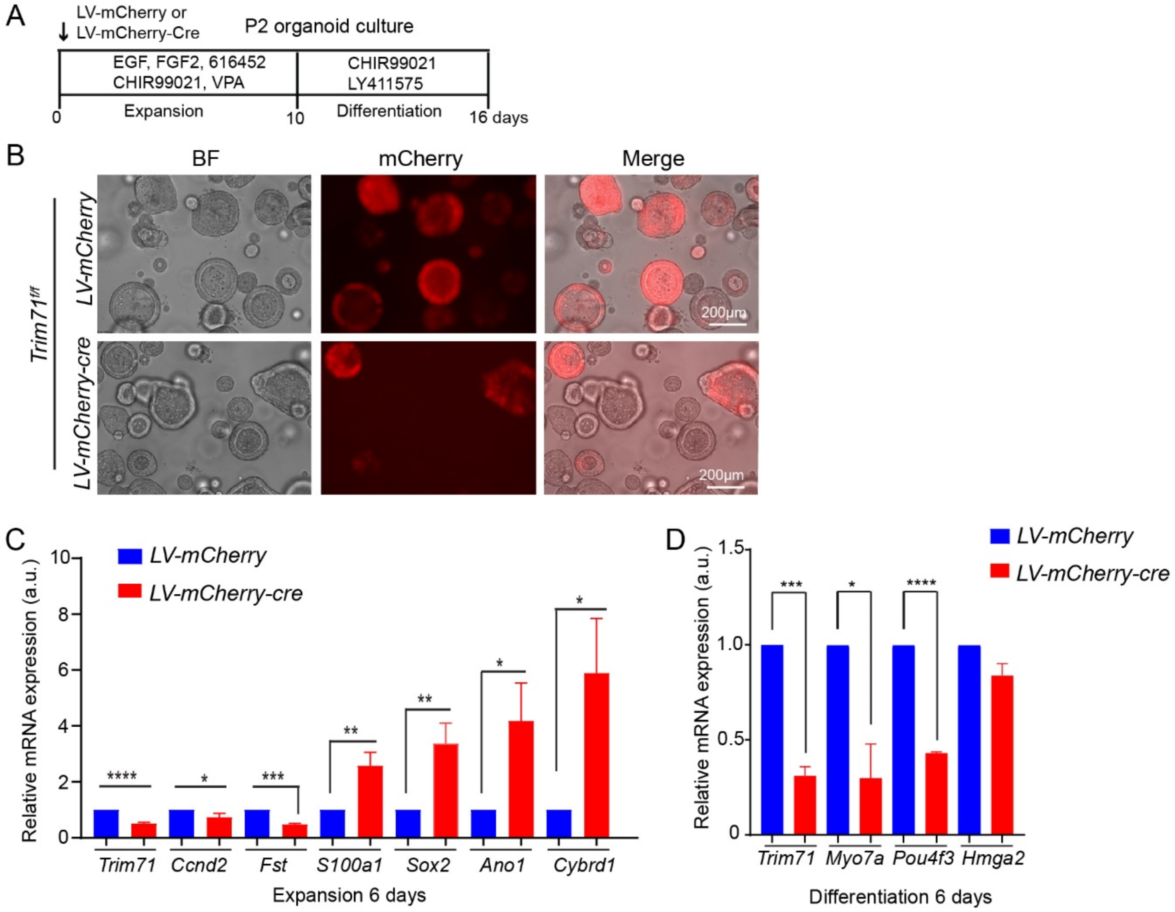


**Supplemental Figure 2. Loss of *Trim71* inhibits supporting cell reprogramming and impedes hair cells formation. (A)** Experimental scheme. **(B)** Bright field and red fluorescent images of cochlear organoid cultures from stage P2 *Trim71* floxed mice infected with lentivirus expressing mCherry and cre, or mCherry only (control). **(C)** RT-qPCR of progenitor (*Trim71, Ccnd2, Fst)* and supporting cell-specific (*S100a1, Ano1, Cybrd1)* mRNA expression in control (LV-mCherry) and *Trim71* deficient (LV-mCherry-cre) organoids after 6 days of expansion (n=3, two independent experiments). **(D)** RT-qPCR of progenitor (*Trim71, Hmga2*) and hair cell-specific (*Myo7a, Pou4f3*) mRNA expression in control (LV-mCherry) and *Trim71* deficient (LV-mCherry-Cre) organoids after 6 days of differentiation (n=3, two independent experiments). Two-tailed, unpaired *t* test was used to calculate *P* values in (C) and (D). *$P \leq 0.05$, **$P < 0.01$, ***$P < 0.001$ and ****$P < 0.0001$.

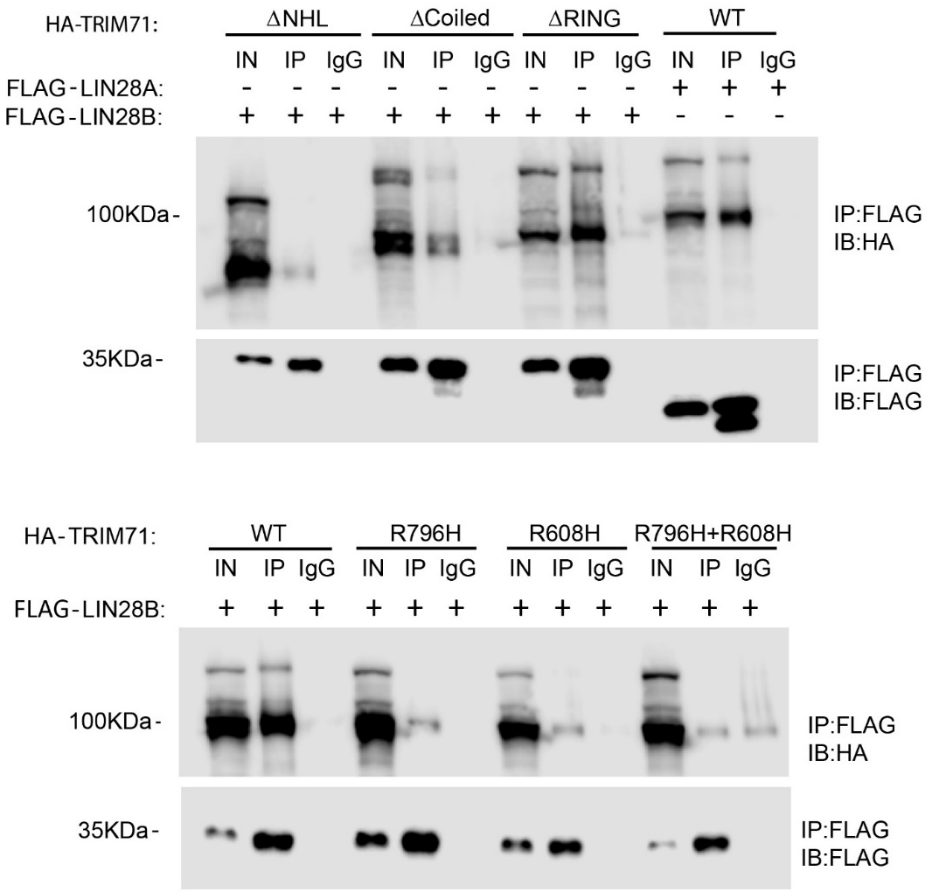


**Supplemental Figure 3. TRIM71's interaction with LIN28B requires a functional NHL domain.** Co-immunoprecipitation (IP) experiments were conducted using HEK293T cells that co-expressed FLAG-tagged LIN28B (or LIN28A) and HA-tagged full-length TRIM71 or mutant TRIM71 proteins. TRIM71 mutant proteins either lacked TRIM71's NHL (ΔNHL), coiled-coil (ΔCoiled) or RING (ΔRING) domain, or carry point mutations in the NHL domain (R796H, R608H, R796H+R608H) that render the domain inactive. Anti-FLAG antibody and IgG antibody (negative control) were used for pull-down.1% of input material (IN) and eluates of pull-down with anti-FLAG antibody (IP) or IgG antibody (IgG) were analyzed by immunoblot (IB) against the HA-tag and FLAG-tag.

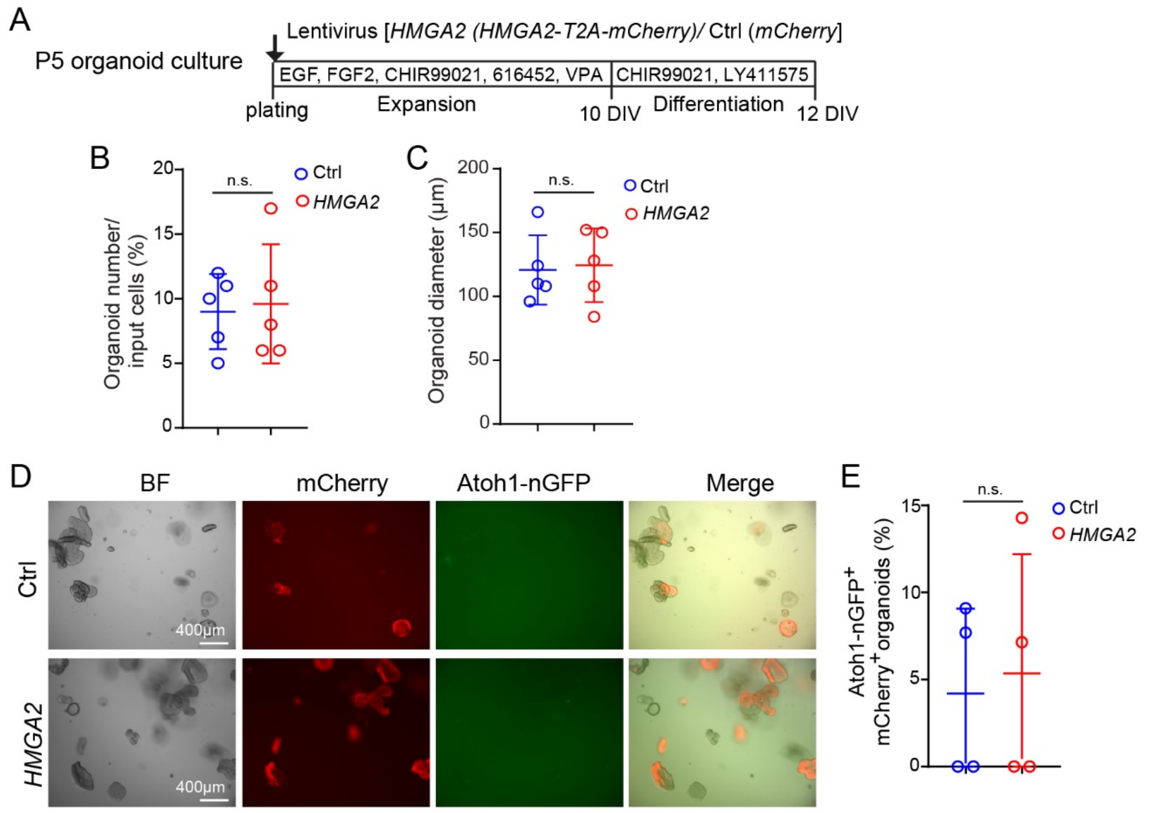


**Supplemental Figure 4. HMGA2 overexpression does not enhance the mitotic or hair cell-forming potential of cochlear supporting cells/Kölliker's cells.** (**A**) Experimental scheme. Organoid cultures were established with cochlear epithelial cells from stage P5 Atoh1-nGFP transgenic mice. (**B**) Colony forming efficiency in *HMGA2* overexpressing and control cultures at 10 DIV. (**C**) Organoid diameter in *HMGA2* overexpressing and control organoid cultures at 10 DIV. (**D**) Representative, low-power bright field (BF) and green fluorescent (Atoh1-nGFP) images of *HMGA2* overexpressing and control organoid cultures. **(K)** Quantification of Atoh1-GFP$^{+}$ organoids in (E). Individual date points represent the average value per animal. Two-tailed, unpaired *t* test was used to calculate *P* values. Abbreviation: n.s., not significant.
