## Supplemental Figures for "Reactivation of the progenitor gene Trim71 enhances the mitotic and hair cell-forming potential of cochlear supporting cells"

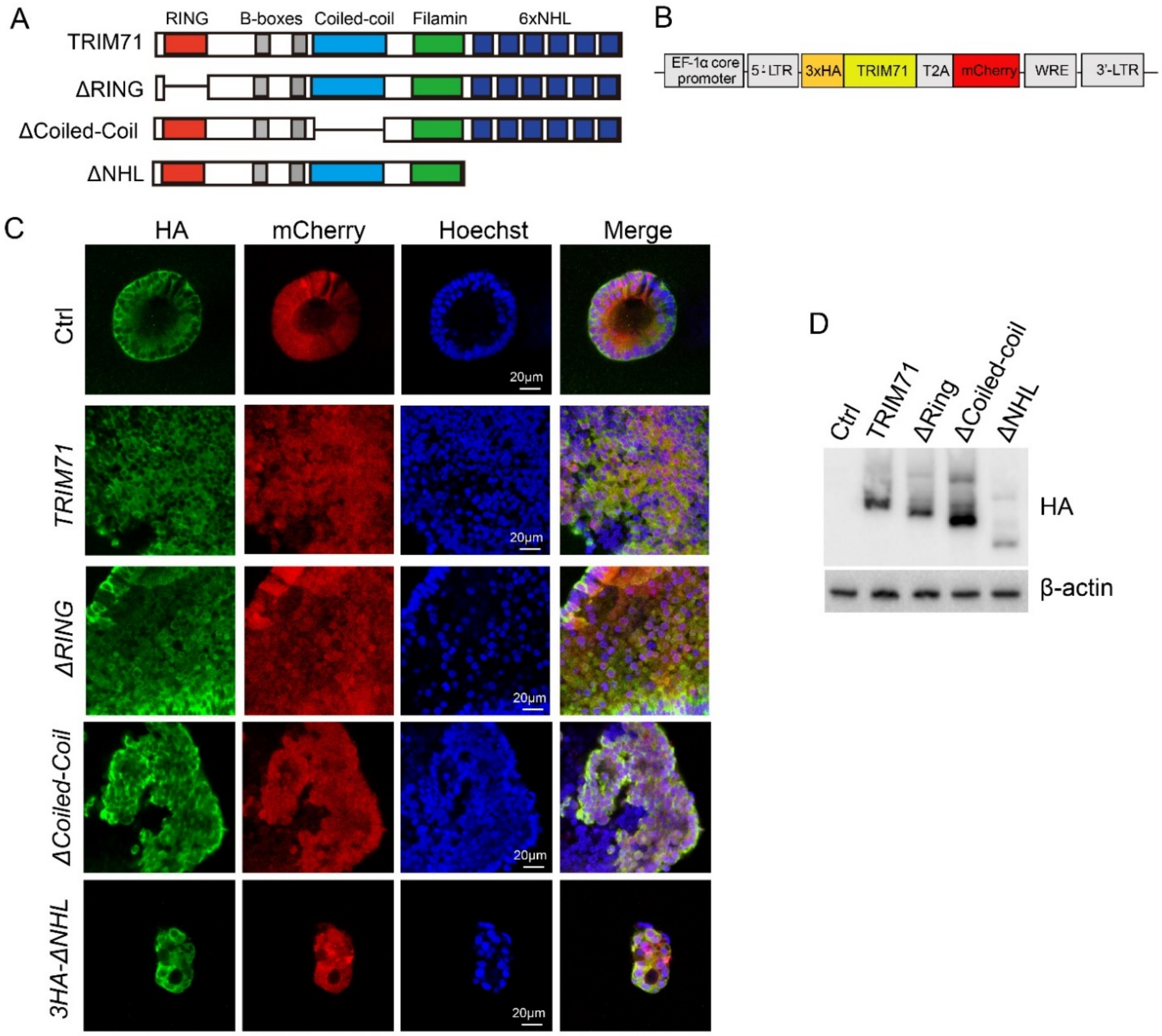


**Supplemental Figure 1. Structure of TRIM71 mutants. (A)** Schematic of human TRIM71 protein and mutant variants that lack the RING (red, ∆ RING), Coiled coil (light blue, ∆ Coiled coil) or NHL domains (dark blue, ∆ NHL). **(B)** Schematic of expression cassette. **(C)** Confocal images of P5 cochlear organoids infected with lentivirus expressing TRIM71 protein and TRIM71 mutant variants. MCherry (red) labels infected cells and HA immunostaining (green) visualizes TRIM71 protein expression. Note control cultures were infected with virus that expresses HA only. **(D)** Protein expression levels of full-length TRIM71 (TRIM71) and TRIM71 mutant variants (△RING, △Coiled-Coil and △NHL) in P5 cochlear organoids at 10 days of expansion. Shown are immunoblots against HA and β-actin.

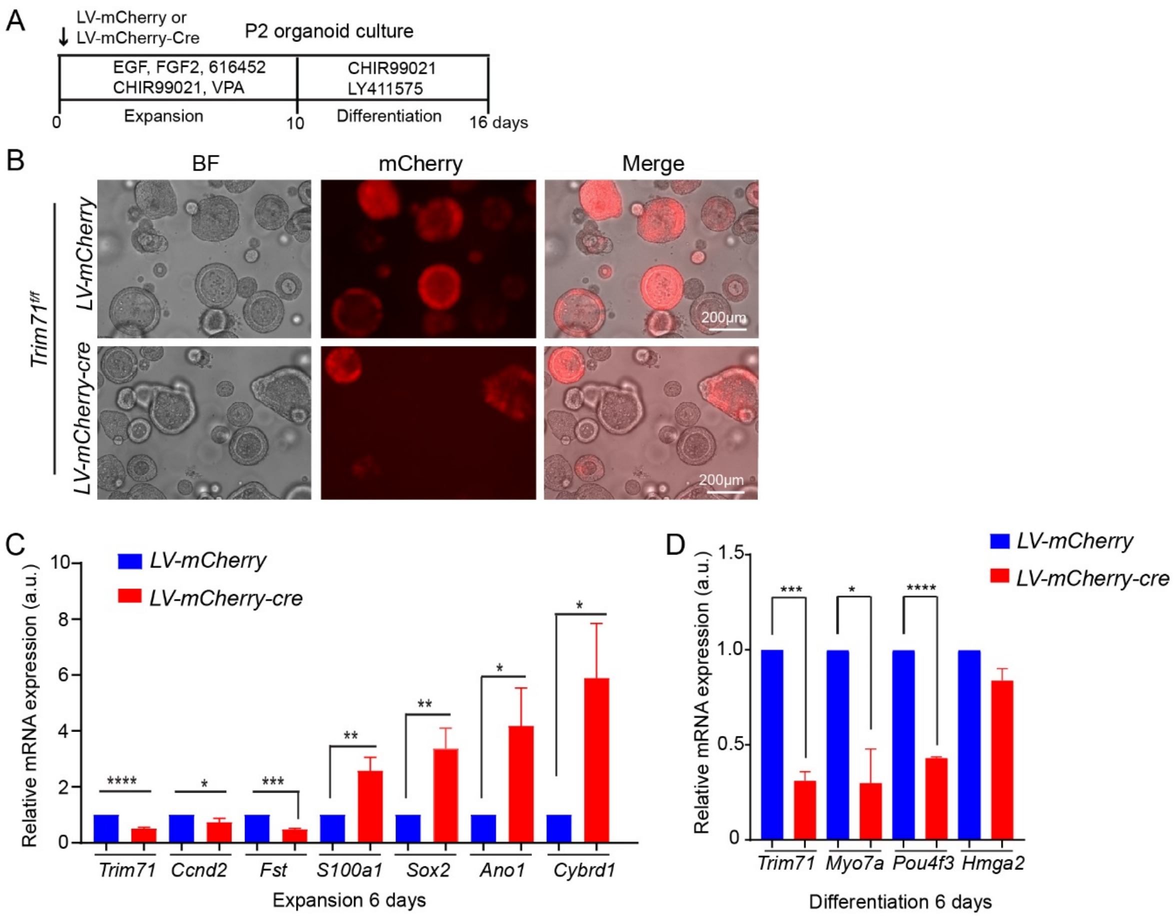


**Supplemental Figure 2. Loss of *Trim71* inhibits supporting cell reprogramming and impedes hair cells formation.** **(A)** Experimental scheme. **(B)** Bright field and red fluorescent images of cochlear organoid cultures from stage P2 *Trim71* floxed mice infected with lentivirus expressing mCherry and cre, or mCherry only (control). **(C)** RT-qPCR of progenitor (*Trim71, Ccnd2, Fst)* and supporting cell-specific (*S100a1, Ano1, Cybrd1)* mRNA expression in control (LV-mCherry) and *Trim71* deficient (LV-mCherry-cre) organoids after 6 days of expansion (n=3, two independent experiments). **(D)** RT-qPCR of progenitor (*Trim71, Hmga2*) and hair cell-specific (*Myo7a, Pou4f3*) mRNA expression in control (LV-mCherry) and *Trim71* deficient (LV-mCherry-Cre) organoids after 6 days of differentiation (n=3, two independent experiments). Two-tailed, unpaired *t* test was used to calculate *P* values in (C) and (D). *$P \leq 0.05$, **$P < 0.01$, ***$P < 0.001$ and ****$P < 0.0001$.

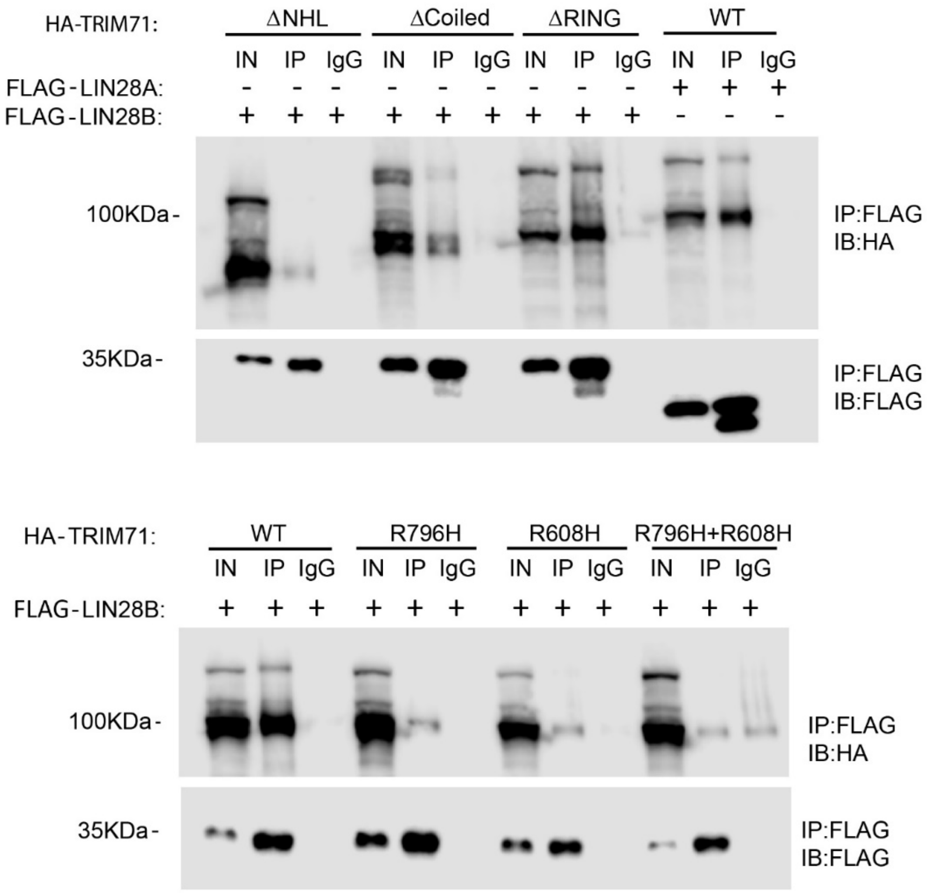


**Supplemental Figure 3. TRIM71's interaction with LIN28B requires a functional NHL domain.** Co-immunoprecipitation (IP) experiments were conducted using HEK293T cells that co-expressed FLAG-tagged LIN28B (or LIN28A) and HA-tagged full-length TRIM71 or mutant TRIM71 proteins. TRIM71 mutant proteins either lacked TRIM71's NHL (ΔNHL), coiled-coil (ΔCoiled) or RING (ΔRING) domain, or carry point mutations in the NHL domain (R796H, R608H, R796H+R608H) that render the domain inactive. Anti-FLAG antibody and IgG antibody (negative control) were used for pull-down.1% of input material (IN) and eluates of pull-down with anti-FLAG antibody (IP) or IgG antibody (IgG) were analyzed by immunoblot (IB) against the HA-tag and FLAG-tag.

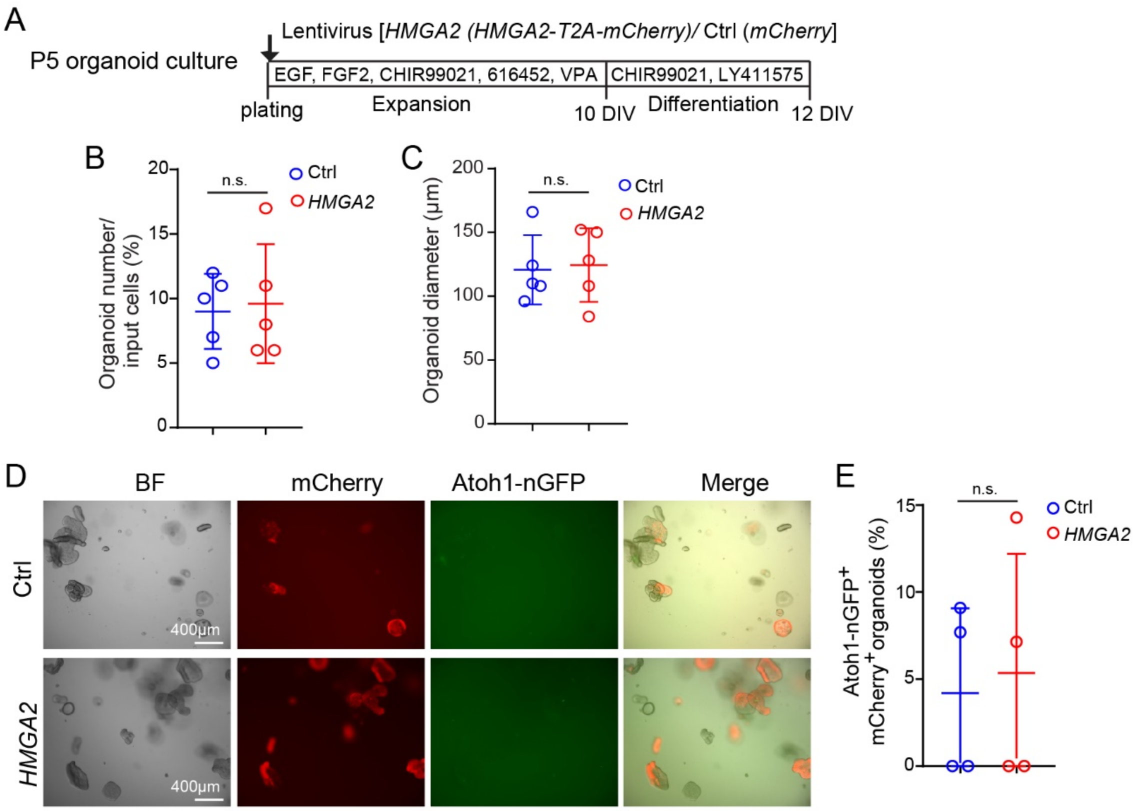


**Supplemental Figure 4. HMGA2 overexpression does not enhance the mitotic or hair cell-forming potential of cochlear supporting cells/Kölliker's cells.** (**A**) Experimental scheme. Organoid cultures were established with cochlear epithelial cells from stage P5 Atoh1-nGFP transgenic mice. (**B**) Colony forming efficiency in *HMGA2* overexpressing and control cultures at 10 DIV. (**C**) Organoid diameter in *HMGA2* overexpressing and control organoid cultures at 10 DIV. (**D**) Representative, low-power bright field (BF) and green fluorescent (Atoh1-nGFP) images of *HMGA2* overexpressing and control organoid cultures. **(K)** Quantification of Atoh1-GFP$^+$ organoids in (E). Individual date points represent the average value per animal. Two-tailed, unpaired *t* test was used to calculate *P* values. Abbreviation: n.s., not significant.
